# MOXD1 is a gate-keeper of organ homeostasis and functions as a tumor-suppressor in neuroblastoma

**DOI:** 10.1101/2023.01.17.524367

**Authors:** Elina Fredlund, Stina Andersson, Elien Hilgert, Guadalupe Álvarez-Hernán, Ezequiel Monferrer, Sinan Karakaya, Tomas Gregor, Siebe Loontiens, Jan Willem Bek, Estelle Lecomte, Emma Magnusson, Enrika Miltenyte, Marie Cabirol, Michail Kyknas, Niklas Engström, Marie Arsenian Henriksson, Emma Hammarlund, Rosa Noguera, Frank Speleman, Johan van Nes, Sofie Mohlin

**Affiliations:** Division of Pediatrics, Department of Clinical Sciences, Lund University; Lund, Sweden; Lund University Cancer Center, Lund University; Lund, Sweden; Lund Stem Cell Center, Lund University; Lund, Sweden; Center for Medical Genetics, Ghent University; Ghent, Belgium; Department of Pathology, Medical School, University of Valencia-INCLIVA Biomedical Health Research Institute; Valencia, Spain; Low Prevalence Tumors, Centro de Investigación Biomédica En Red de Cáncer (CIBERONC), Instituto de Salud Carlos III; Madrid, Spain; Department of Microbiology, Tumor and Cell Biology (MTC), Biomedicum B7, Karolinska Institute; Stockholm, Sweden; Department of Laboratory Medicine, Lund University; Lund, Sweden; Department of Oncogenomics, Academic Medical Center, University of Amsterdam; Amsterdam, the Netherlands

## Abstract

Neuroblastoma is a childhood cancer believed to result from dysfunctional development. Its origin during embryogenesis remains poorly understood. The lack of appropriate models has hindered in-depth mapping of tumor-driving events. Here, we identify a novel tumor-suppressor gene that predicts poor survival in high-risk disease, by applying bulk and single cell RNA sequencing data of neuroblastoma and human fetal adrenal glands. Trunk neural crest-specific MOXD1 discriminates cell populations during normal and tumor development, with implications for deciphering neuroblastoma cell origin. We created an embryonic conditional knockout model and show that cell type-specific loss of *MOXD1* leads to disrupted organ homeostasis and failed adrenal gland formation, home for neuroblastoma. We show that MOXD1 is a tumor suppressor gene in zebrafish, chick, and mice *in vivo* models.

**One-Sentence Summary:** Neural crest-specific MOXD1 is a *de novo* tumor-suppressor gene in childhood cancers arising during embryogenesis.

---

Neuroblastoma (NB) is the most common cause of cancer-related deaths in infants, and most patients have received their diagnosis before the age of five (*1*). NB aggressiveness is influenced by a number of factors, including loss of 6q. Patients present with a five-year survival rate of approximately 90-95 % vs. 40-50 % for low- and high-risk NB, respectively (*2*, *3*). Two distinct cellular phenotypes are suggested to foster NB development and progression: lineage-committed adrenergic (ADRN) and immature mesenchymal (MES) (*4*). Characterization of the two phenotypes is based on transcriptomic and epigenetic profiling, and ADRN cells have been found to grow more aggressively in mouse models *in vivo* while MES cells are more resistant to treatment with chemotherapy and ALK inhibitors, pinpointing intratumor heterogeneity and plasticity (*4*).

Most NBs are found in the abdominal area and the primary site of tumors is commonly related to the adrenal glands or the sympathetic ganglia (*3*, *5*, *6*). NBs originate from a neural crest (NC) progenitor population during early embryogenesis. The multipotent NC cells (NCC) undergo epithelial-to-mesenchymal transition (EMT) to initiate migration and eventually differentiate into a plethora of cell types. Cells that derive from the trunk derivative of the NC give rise to Schwann cell precursors (SCPs), melanocytes, cells of the sympathetic nervous system and chromaffin cells in the adrenal medulla. NB is believed to form following disrupted trunk NC (tNC) cell (tNCC) differentiation; however, the NB founder cell type is still unknown. Lately, many efforts have been made to map the cellular composition of normal developing human and murine adrenal medulla with single cell RNA sequencing (scRNA seq) to the features of NB cells (*7*–*18*). Findings show that NB cells relate to various stages during normal neuroblast maturation. However, discrepancies in interpretation of data, the usage of different species and developmental time points, and analyses restricted to subtypes of NB warrants further studies. So far, there is no classification of NBs as defined by their cellular origin.

Monooxygenase DBH-like 1 (MOXD1) is a tNC-enriched gene as demonstrated by *in situ hybridization* and bulk RNAseq of sorted NCCs during avian development (*19*, *20*). MOXD1 is mainly localized in the rough endoplasmic reticulum (*21*, *22*), and homologs to human *MOXD1* gene has been found in zebrafish, chicken, and mouse (62%, 77% and 84% identity, respectively). MOXD1 is a member of the copper monooxygenase family that requires copper ions to carry out the incorporation of hydroxyl groups into its substrates. The well characterized protein dopamine beta-hydroxylate (DBH) is also part of the copper monooxygenase family (*22*, *23*). While many of the essential functions carried out by DBH in chromaffin cells and neurons of the central nervous system are known, the function of MOXD1 has not yet been determined.

MOXD1 emerges in EMT-transformed and migrating tNCCs, and connections between tNCCs and NB initiation during specific embryonic time windows have been suggested. Thus, here we investigated the impact of MOXD1 on neural crest and NB development. We show that MOXD1 is a novel tumor suppressor gene in NB, using patient material and clinical data sets, *in vivo* models from three different animal species, and biologically functional assays. MOXD1 discriminates between the ADRN and MES cell phenotypes, with cell-specific expression in the NC-like MES cells. In line with this, MOXD1 is restricted to SCPs during normal development of human and mouse adrenal medulla, reflecting the overlap between SCPs and MES NB cell phenotypes. We created a conditional embryonic knockout model with tNCC-specific loss of MOXD1, leading to disrupted tissue architecture and altered organogenic fate. Our findings depict a *de novo* tumor suppressor gene and potentiates identification of subtype-specific cells-of-origin in NB.

## Results

### Loss of MOXD1 predicts worse outcome in high-risk neuroblastoma patients

To analyze the clinical impact of MOXD1 in neural crest-derived NB, we utilized two independent patient cohorts consisting of NBs from all risk groups. Stratifying patients from the SEQC cohort (*24*) (*n* = 498) based on their *MOXD1* expression showed that tumors with the lowest (1^st^ quartile) expression of *MOXD1* present with worse overall- and event free survival (Fig. 1A and Supplementary Fig. S1A). *MOXD1* was an independent predictor of survival also after correcting for common risk factors age at diagnosis, INSS (International Neuroblastoma Staging System) stage, and *MYCN* amplification by multivariate Cox regression (Fig. 1B; Supplementary Fig. S1B). These results were confirmed in a second cohort (Kocak (*26*), *n* = 649; Fig. 1C and Supplementary Fig. S1B-C). Using the same two patient cohorts, we analyzed *MOXD1* expression according to INSS stages and low- and high-risk groups, and observed a negative correlation between *MOXD1* and more advanced tumor stage and high-risk NB (Fig. 1D-E and Supplementary Fig. S1D). Expression of MOXD1 increased in Stage 4s as compared to Stage 4, which is coherent with spontaneous regression of Stage 4s tumors (Fig. 1E and Supplementary Fig. S1D). Ganglioneuromas (GNs) are composed of abnormally growing and migrating immature neuroblasts, and classify as neuroblastic tumor together with ganglioneuroblastoma and NB. However, while NB is highly malignant, GNs are benign tumors, often asymptomatic. Sequencing data from patients with either of these two forms (*27*) showed significantly lower expression of *MOXD1* in NB as compared to GN (Fig. 1F), being one of the top 20 downregulated genes (*27*). We examined *MOXD1* mutational status and possible translocations in 87 NBs of all stages but found no such alterations. Copy-number profiles of the same 87 NBs however indicated presence of chromosomal losses surrounding the *MOXD1* locus (6q23.2; Fig. 1G). The group of high-risk NB patients with loss of distal 6q (32/542; 5.9 %) present with extremely low survival probability (*28*). Applying the cohort from Depuydt *et al* (2018) consisting of exclusively high-risk tumors (Depuydt, *n* = 556), we found a small, yet distinct group of tumors presenting with loss of *MOXD1* (Fig. 1H). We confirmed that expression of *MOXD1* was low in patients with loss of 6q in this cohort (Supplementary Fig. S2A). Patients with chromosomal loss surrounding the *MOXD1* locus presented with worse outcome when comparing to patients with normal chromosome 6q karyotype (i.e., no chromosomal aberrations surrounding the MOXD1 locus; patients with loss of 6q but not MOXD1 were excluded), noteworthy in this patient group already presenting with dismal prognosis (5-year survival rate of <20% as compared to ~40% in patients without any gains or losses in this region; Fig. 1I; Supplementary Fig. S2B). There was no difference in outcome when comparing normal 6q karyotype vs MOXD1 gain or gain/losses collectively (Supplementary Fig. S2C-F). Together, this demonstrates that low expression or loss of *MOXD1* correlates to unfavorable disease in NB.

**Fig. 1.**
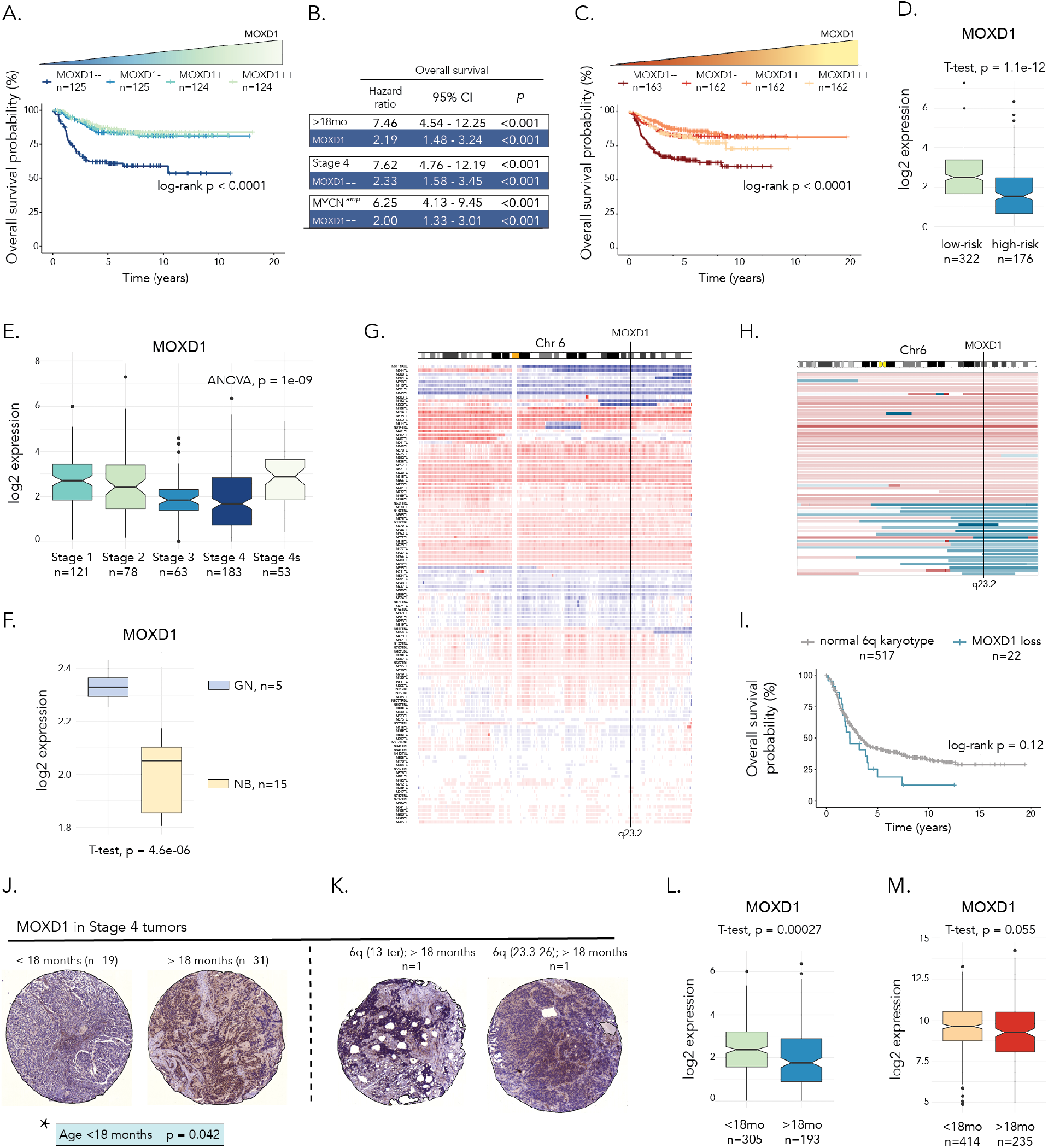
Low *MOXD1* expression predicts worse prognosis in neuroblastoma patients. **(A**) Kaplan-Meier survival curves with log-rank *p*-value for NB patients in the SEQC cohort (*n*=498) stratified into four groups (quartiles) based on *MOXD1* mRNA expression. **(B)** Prognostic effect of *MOXD1* mRNA expression for NB patients (SEQC cohort, 1^st^ vs 2^nd^-4^th^ quartile of *MOXD1* expression) with adjustment for age at diagnosis, INSS stage of disease and *MYCN* amplification status, respectively. Hazard ratios, 95% confidence intervals (CI) and log rank *p*-values are given by multivariate Cox regression analyses. **(C)** Kaplan-Meier survival curves with log-rank *p*-value for NB patients in the Kocak cohort (*n*=649) stratified into four groups (quartiles) based on *MOXD1* mRNA expression. (**D-E**) *MOXD1* expression in NB patients (SEQC cohort) stratified according to low- or high risk (D) or INSS stages (E). Number of patients and *p*-value by t-test and ANOVA as indicated. (**F**) *MOXD1* expression in NB vs ganglioneuroma (GN) patients (cohorts from Tao *et al* (2020)). Number of patients and *p*-value by t-test as indicated. (**G**) DNA copy-number aberrations on chromosome 6 from 87 NBs of all stages. Gains, red; losses, blue. (**H**) Detailed view of gains (red) and losses (blue) on chromosome 6 from high-risk patients in Depuydt cohort (*n*=556). The black line indicates the chromosomal location of MOXD1 (6q23.2). Maximum size of aberration was set to 180 Mb. (**I**) Kaplan-Meier plot of overall survival of high-risk NB patients (Depuydt; *n* = 541). Patients stratified by loss of *MOXD1* or no loss of *MOXD1*. Patients with distal 6q mutations not affecting *MOXD1* and patients lacking overall survival information were excluded. *p*-value by log-rank test as indicated. (**J-K**) Representative images of tumor cores stained for MOXD1 in different age groups (≤ 18 months vs > 18 months at diagnosis (J)) and tumors with loss of 6q (K). *Statistical analysis of significant correlations between MOXD1 expression and age at diagnosis. (**L-M**) *MOXD1* expression in NB patients (SEQC (L) and Kocak (M) cohorts) stratified according to age at diagnosis (<18 months vs >18 months). *p*-value by t-test as indicated.

### The prognostic role of MOXD1 is tumor-type specific

Comparing NB to malignant melanoma, another tNC-derived cancer, using data from the Cancer Cell Line Encyclopedia cohort (*29*), showed that *MOXD1* expression was significantly lower in NB (Supplementary Fig. S3A). To investigate whether MOXD1 could still have a prognostic role in malignant melanoma, we compared expression in primary (n=16) and metastatic (n=188) tumors (*30*), and found a small increase in expression in metastatic tumors (Supplementary Fig. S3B). In the non-tNC derived tumor forms colorectal (*31*) and breast cancer (*32*) we compared *MOXD1* expression in Stage 1/2/3 vs. Stage 4 and localized vs. metastatic, respectively. There were no distinct differences in expression between these groups (Supplementary Fig. S3C-D).

### MOXD1 protein expression correlates with age in stage 4 tumors

We stained a tissue microarray (TMA) consisting of material from 50 patients restricted to Stage 4 and MYCN-amplified NBs by immunohistochemistry (Fig 1J). Two cases presented with loss of distal 6q (Fig 1K). Albeit few patients (2/50; 4%), this is in line with the published proportion of tumors with loss of distal 6q among the high-risk NB patient cohort (*28*). We observed a heterogenous expression pattern of MOXD1 across all tumors, and the higher MOXD1 expression, the higher was the variation in intratumor intensity. While there was no correlation to outcome within this patient group with already poor prognosis, expression of MOXD1 protein correlated to age at diagnosis, with lower heterogeneity of MOXD1 in children <18 months (Fig. 1J). Tumors with loss of 6q presented with ~30% positive cells each, but only 1% and 4% of those were stained with high intensity. These cores were thus not depleted of MOXD1, highlighting intra-tumor heterogeneity and the presence of surrounding non-tumor cells in these tissues. Both cases with loss of 6q were patients >18 months (Fig. 1K). We went back to the two data sets analyzed in Fig. 1A and 1C (SEQC and Kocak), distributed to reflect the clinical situation with patients from all stages and risk groups, and found that *MOXD1* mRNA expression was lower in patients over the age of 18 months at diagnosis (Fig. 1L-M).

### MOXD1 is restricted to undifferentiated MES type cells in neuroblastoma

Analysis of scRNAseq data from 11 NBs (*7*) showed enrichment of *MOXD1* in the clusters with undifferentiated tumor cells (Fig. 2A). MES and ADRN NB cells display distinct genetic phenotypes; undifferentiated NC-like, and committed adrenergic, respectively. Expression analysis of NB cell lines and healthy NC cells showed that NC and MES cells had higher *MOXD1* levels as compared to ADRN cells that had virtually absent expression of this gene (Fig. 2B), in line with the data from primary patient material (Fig. 2A). We further analyzed sequencing data from three isogenic MES and ADRN cell line pairs. The 691 and the 717 pairs are primary cell cultures while the SH-EP2 (MES phenotype) and SH-SY5Y (ADRN phenotype) subclones stem from the conventional NB cell line SK-N-SH. All three MES cell lines expressed *MOXD1* while expression was absent in corresponding ADRN cells (Fig. 2C). Analysis of the 691-MES and 691-ADRN cells *in vitro* confirmed that *MOXD1* expression was restricted to MES cells (Fig. 2D).

**Fig. 2.**
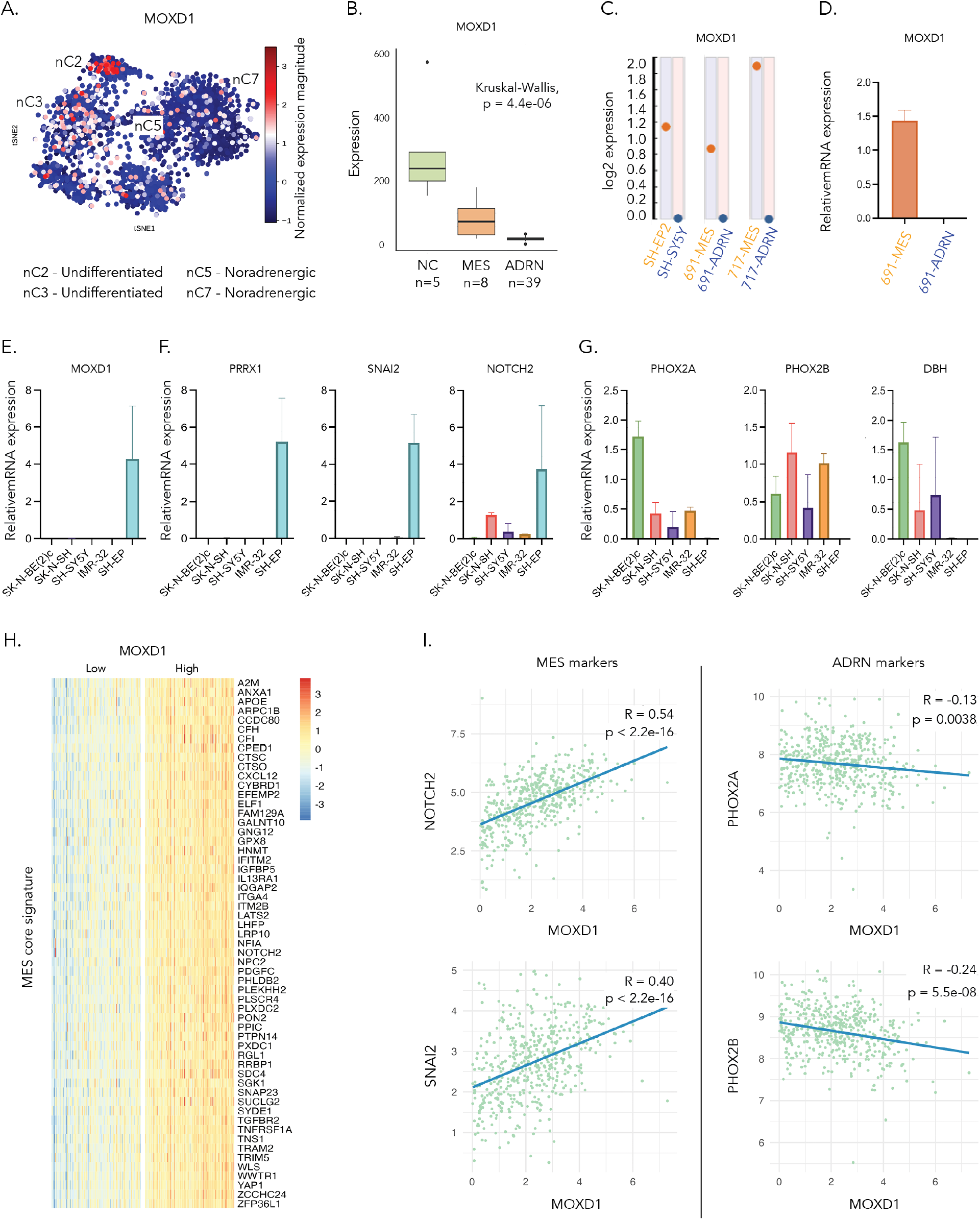
*MOXD1* is expressed in MES-like NB cells. **(A**) Mapping of *MOXD1* mRNA expression in the t-distributed stochastic neighbor embedding (tSNE) of human NB single nuclei data from Bedoya-Reina *et al* (2021), analyzed using PAGODA. nCx (2, 3, 5 and 7) are selected clusters from published data above. **(B**) *MOXD1* mRNA expression in NB cells with ADRN and MES gene signatures and in normal neural crest (NC). Number of samples (*n*) for each cell and *p*-value by the Kruskal-Wallis test as indicated. (**C**) RNA sequencing derived log2 transformed expression of *MOXD1* in isogenic NB cell line pairs. (**D**) *MOXD1* mRNA expression in patient-derived 691-cells of MES and ADRN subtypes *in vitro*, as determined by quantitative real-time PCR (qPCR). Bars represent standard deviation, *n* = 3 biologically independent repeats. (**E**) *MOXD1* mRNA expression in three NB cell lines with ADRN phenotype (SK-N-BE(2)c, SH-SY5Y, and IMR-32), one with mixed phenotype (SK-N-SH), and one with MES phenotype (SH-EP) as assessed by qPCR. Error bars denote standard deviation from *n* = 2-3 biologically independent repeats per cell line. (**F-G**) Expression of MES-(F) and ADRN-(G) associated genes as assessed by qPCR. *n* = 3 biologically independent repeats. **(H)** Heatmap of the genes that were considered among the 500 most DEGs between the 1^st^ and 4^th^ quartile of *MOXD1* expression in the SEQC cohort and also included in the MES and ADRN core signatures from van Groningen *et al* (2017). (**I**) Correlation between *MOXD1* mRNA expression and two mesenchymal (*NOTCH2* and *SNAI2*), or two adrenergic (*PHOX2A* and *PHOX2B*) markers. Pearson correlation coefficients (R) and two-sided *p*-values. Coefficients of determination and *p*-values generated by linear regression as indicated.

In concordance with isogenic cell pairs (*4*), *MOXD1* and MES-associated genes *PRRX1*, *SNAI2* and *NOTCH2*(*4*) were restricted to the MES-like SH-EP cells (Fig. 2E-F). The three ADRN-like cell lines SK-N-BE(2)c, SH-SY5Y, and IMR-32, and the mixed ADRN/MES cell line SK-N-SH (*33*, *34*) were devoid of *MOXD1* and MES-associated markers, but instead expressed ADRN-associated genes *PHOX2A*, *PHOX2B*, and *DBH* (Fig. 2G). We separated the SEQC cohort (analyzed in Fig. 1) into *MOXD1*^Low^ (1^st^ quartile) and *MOXD1*^High^ (4^th^ quartile) groups and observed that the MES core signature (*4*) was highly enriched in the *MOXD1*^High^ tumors (Fig. 2H). *MOXD1* correlated positively with MES-, and negatively with ADRN-phenotype markers (Fig. 2I).

### MOXD1 is enriched in Schwann cell precursor cells

Analysis of scRNAseq data from E9.5 – E12.5 mouse embryos showed enrichment of *MOXD1* in tNCCs and SCPs, specifically (Fig. 3A). Expression data from human embryonic stem cells (ESCs) and tNC derived from these ESCs by *in vitro* differentiation (*35*) also demonstrated *MOXD1* expression specifically in tNCCs (Fig. 3B). Further analyses of scRNAseq data from precursor cells of the sympathoadrenal lineage during mouse (*36*) and human (*14*) development demonstrated that *MOXD1* expression was, in alignment with above, highly enriched in SCPs (Fig. 3C-E). It has been shown that NB MES-like cells cluster together with SCPs (*37*), and these data demonstrate a cell type-specific expression of *MOXD1* during normal development in both mouse and human.

**Fig. 3.**
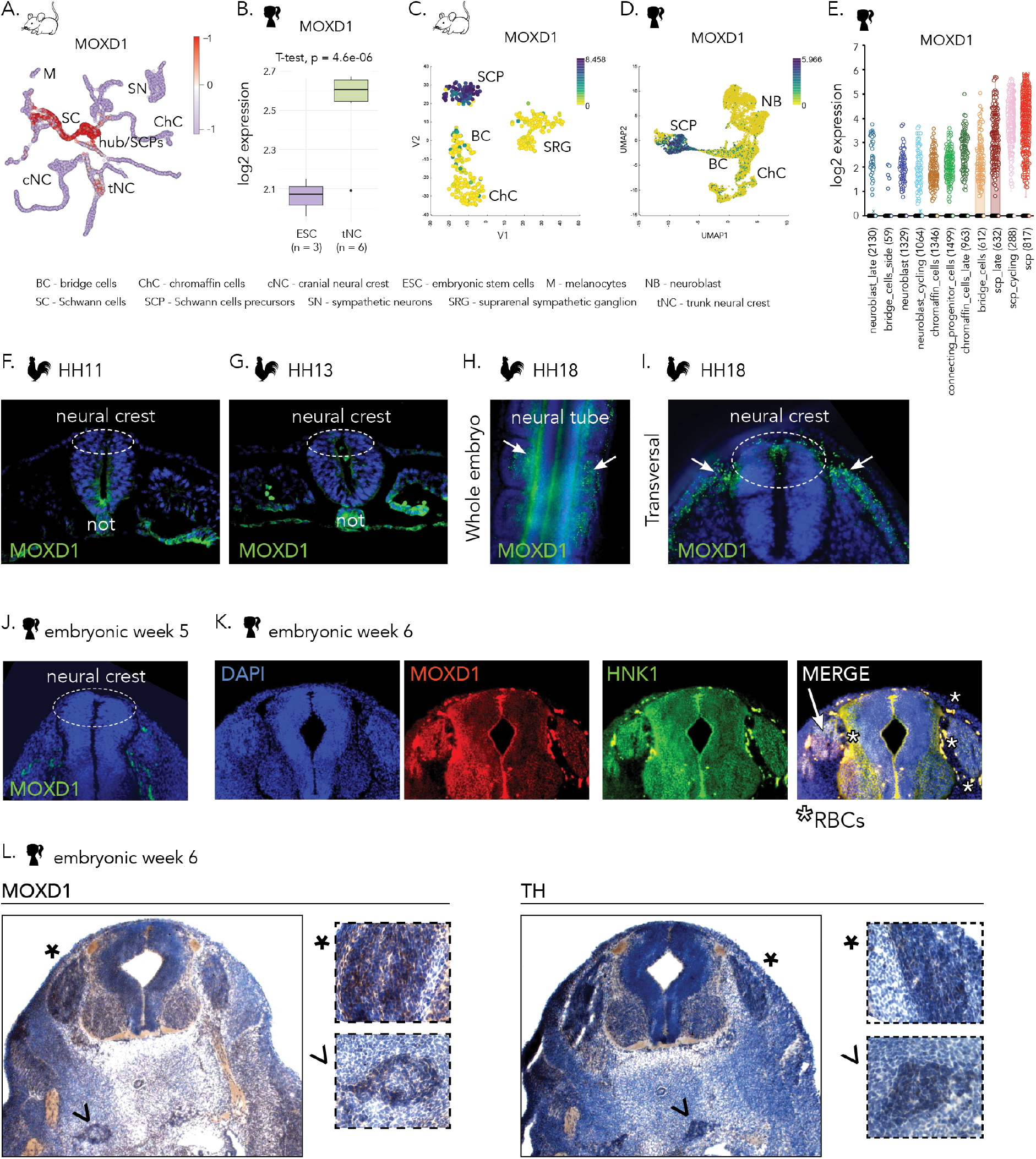
*MOXD1* is expressed in trunk neural crest and Schwann cell precursors during normal adrenal gland development. **(A)** Mapping of *MOXD1* mRNA expression in the *Uniform Manifold Approximation and Projection* (UMAP) of single cells from mouse neural crest and Schwann cell linages from Kastriti *et al* (2022), analyzed using PAGODA. **(B)** Expression of *MOXD1* mRNA in human embryonic stem cells (ESC) as compared to *in vitro* ES cell-derived trunk neural crest using data from Frith *et al* (2018). **(C)** Visualization of *MOXD1* by cell type clustering on single cell analysis of the mouse neural crest from Furlan *et al* (2017). Color scale shows log2 transformed expression. **(D-E)** Visualization of *MOXD1* by cell type clustering on single cell analysis of the human adrenal lineage from Jansky *et al* (2021) (D). Color scale shows log2 transformed expression. Expression in individual cell types of the adrenal lineage is indicated by color (E). **(F-G)** Immunofluorescence staining of MOXD1 protein in transversal sections of the trunk axial level of chick embryonic body of trunk neural crest pre-migratory stages HH11 (F) and HH13 (G). DAPI (blue) is used to visualize nuclei. **(H-I)** Immunofluorescence staining of MOXD1 protein in whole embryo view (H) and transversal sections (I) of the trunk axial level of chick embryonic body of trunk neural crest cell at post-migratory stage HH18. DAPI (blue) is used to visualize nuclei. **(J)** Immunofluorescence staining of MOXD1 protein in section of the trunk axial level of human embryonic body, of trunk neural crest cell at pre-migratory stage embryonic week (ew) 5. DAPI (blue) is used to visualize nuclei. **(K)** Immunofluorescent staining of MOXD1 and HNK1 (to identify migrating neural crest cells) protein in sections of the trunk axial level of human embryonic body of trunk neural crest cell at post-migratory stage embryonic week 6. DAPI (blue) is used to visualize nuclei. Asterisk indicate auto-fluorescent red blood cells (RBCs; erythrocytes). Arrow indicates true staining. **(L)** Immunohistochemical staining of MOXD1 and TH (to identify sympathetic ganglia and adrenal gland) protein in sections of the trunk axial level of human embryonic body of trunk neural crest cells at post-migratory stage embryonic week 6. Nuclei are counterstained by Hematoxylin&Eosin. * and ^ respectively denote where each magnification is localized in the overview images.

### MOXD1 protein is expressed in human and chick embryos

Thus far, expression of *MOXD1* has only been demonstrated at RNA level in healthy embryos (*19*, *20*) and adult tissues (*21*, *22*, *38*), and by using bulk- and scRNAseq here (Figs. 3A-E). We therefore stained avian and human embryos at different developmental time points to confirm cell- and stage-specific protein expression. Embryos were sectioned at the trunk level of the embryonic body, ensuring trunk-restricted analysis. MOXD1 protein was not detected at pre-migratory NC stages (Fig. 3F-G), while, in conjunction with RNA data, strongly expressed in migrating NCCs (Fig. 3H-I) in chick embryos. These expression patterns were reflected in human embryos at corresponding stages, embryonic weeks ew5 (pre-migratory) and ew6 (migratory), as determined by location and co-staining with migrating neural crest-marker HNK1 (Fig. 3J-L).

### Overexpression of MOXD1 prolongs survival and reduces tumor burden in vivo

To assess the role of *MOXD1* in tumor development, we selected three cell lines with a dominant ADRN phenotype (SK-N-BE(2)c, SK-N-SH, and 691-ADRN), lacking expression of endogenous MOXD1 (Fig. 2E), and used a lentiviral system for sustained overexpression of *MOXD1* (Fig. 4A-J). Overexpression was verified by qPCR and immunofluorescent staining (Fig. 4A-B, E, H). Transduced cells (with MOXD1 overexpression and their control counterpart) were subcutaneously injected into mice to study tumor formation and survival *in vivo*. Mice with injected SK-N-BE(2)c^MOXD1-OE^ cells presented with delayed tumor formation with a mean of 15 days (95% CI 12.8 to 16.8) for tumors to reach a volume of 200mm^3^ as compared to 9 days (95% CI 4.9 to 14.0) for mice injected with control cells (Fig. 4C). The delay in tumor formation was reflected by prolonged survival (Fig. 4D). Injected SK-N-SH^MOXD1-OE^ cells resulted in fewer mice with tumors and prolonged survival (Fig. 4F-G). In addition, injection of 691-ADRN^MOXD1-OE^ cells resulted in prolonged survival, and notably 67% tumor-free mice at endpoint (365 days) as compared to the CTRL group where all mice carried tumors (Fig. 4I-J). Hence, our *in vivo* experiments demonstrate that MOXD1 suppresses NB tumorigenesis.

**Fig. 4.**
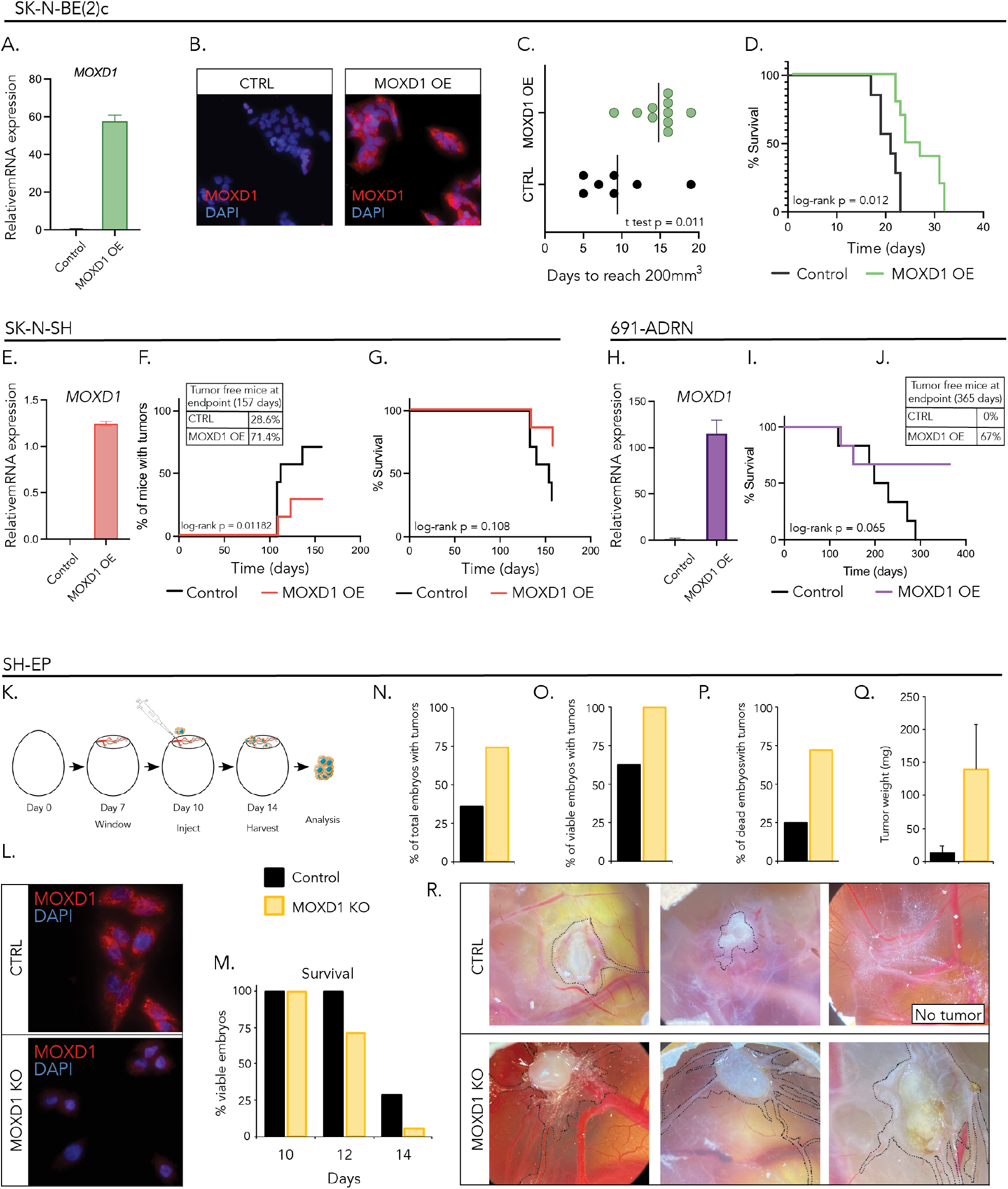
MOXD1 overexpression in ADRN cells prolongs survival *in vivo*. **(A-B)** Confirmation of MOXD1 overexpression by lentiviral transduction at mRNA (A) and protein (B) level in neuroblastoma SK-N-BE(2)c cells by qPCR and immunofluorescence, respectively. DAPI was used to counterstain nuclei. Error bars denote standard deviation from *n* = 3 biologically independent repeats. **(C)** Time to tumor formation in mice subcutaneously injected with control (CTRL) or MOXD1 overexpression (OE) SK-N-BE(2)c cells determined by days to reach a volume of 200 mm^3^. Vertical line indicates mean and the *p*-value by students *t*-test as indicated (*n* = 10 in MOXD1 OE and *n* =7 in CTRL group). **(D)** Kaplan-Meier plot of overall survival in mice of indicated subgroup of SK-N-BE(2)c (*n* = 10 MOXD1 OE, *n* = 7 Control). *p*-value by the log-rank (Mantel Cox) test as indicated. **(E)** Confirmation of MOXD1 overexpression by lentiviral transduction at mRNA level in neuroblastoma SK-N-SH cells by qPCR. Error bars denote standard deviation from *n* = 3 biologically independent repeats. **(F)** Percentage of mice with tumors from indicated subgroup of SK-N-SH cells (*n* = 7 per group). *p*-value by log-rank (Mantel Cox) test. Percentage of tumor-free mice for each subgroup at day 157, the day of experiment termination presented in table format. **(G)** Kaplan-Meier plot of overall survival in mice of indicated subgroup of SK-N-SH (*n* = 7 per group). *p*-value by the log-rank (Mantel Cox) test as indicated. **(H)** Confirmation of *MOXD1* overexpression at mRNA level in NB 691-ADRN cells by qPCR. Error bars denote standard deviation from *n* = 3 biologically independent repeats. **(I)** Kaplan-Meier plot of survival in mice of indicated subgroup of 691-ADRN cells (*n* = 6 per group). *p*-value by the log-rank (Mantel Cox) test as indicated. **(J)** Percentage of tumor free mice at experiment endpoint (365 days). **(K)** Schematic description of the chorioallantoic membrane (CAM) assay. **(L)** Confirmation of CRISPR/Cas9-mediated knockout (KO) of MOXD1 at protein level in neuroblastoma SH-EP cells by immunofluorescence. DAPI was used to counterstain nuclei. **(M)** Survival of chick embryos as presented by percentage of viable embryos at indicated time points. Eggs implanted with CTRL cells are marked in black and *MOXD1* KO in yellow. There are no error bars due to absolute numbers. Implanted eggs; *n*=28 CTRL and *n*=38 MOXD1 KO. **(N)** Number of eggs with detectable tumors at the CAM as presented by percentage at D14. There are no error bars due to absolute numbers. Implanted eggs; *n*=28 CTRL and *n*=38 *MOXD1* KO. **(O)** Number of eggs with viable chick embryos and detectable tumors at the CAM as presented by percentage at D14. There are no error bars due to absolute numbers. Viable embryos in total at D14, *n*=8 CTRL and *n*=2 *MOXD1* KO. **(P)** Number of eggs with dead chick embryos and detectable tumors at the CAM as presented by percentage at D14. There are no error bars due to absolute numbers. Dead embryos in total at D14, *n*=20 CTRL and *n*=25 *MOXD1* KO. **(Q)** Weight of dissected tumors in mg. Error bars indicate standard error of the mean and *p*-value determined by ANOVA. Note that there are few eggs involved in this analysis due to the high number of dead embryos and spread tumors that could not be completely dissected. Weighed tumors at D14, *n*=5 CTRL and *n*=2 *MOXD1* KO. **(R)** Representative images of the CAM in respective group.

### Knockout of MOXD1 enhance tumor burden in vivo

Next, we knocked out MOXD1 by using CRISPR/Cas9 in MES-like SH-EP cells that express high levels of the endogenous gene (Fig. 2E). SH-EP^CTRL^ and SH-EP^MOXD1-KO^ cells were implanted onto the chorioallantoic membrane (CAM) in fertilized eggs *in vivo* (Fig. 4K). For consistency, we implanted at the connecting point of the two largest blood vessels. Knockout of MOXD1 was verified by immunofluorescence (Fig. 4L). To measure the impact on tumorigenesis on embryo viability, we assessed survival after two- and four-days post-implantation (embryonic day E12 and E14, respectively). Implantation of SH-EP^CTRL^ cells did not affect survival until after four days, where viability decreased to 29% (Fig. 4M). In contrast, implantation of SH-EP^MOXD1-KO^ cells reduced survival to 71% already after two days of tumor growth, and to as low as 5% after four days (Fig. 4M). A substantially higher number of tumors was formed with SH-EP^MOXD1-KO^ cells when assessing both viable and dead embryos four days post-implantation (Fig. 4N-P). Knockout of MOXD1 resulted in larger tumors (Fig. 4Q). As visualized before tumor dissection, SH-EP^MOXD1-KO^ cells were also more migratory (Fig. 4R).

### MOXD1 overexpression affects colony growth in a three-dimensional setting

We further examined effects of *MOXD1* overexpression *in vitro*. SK-N-BE(2)c^CTRL^ and SK-N-BE(2)c^MOXD1-OE^ cells were cultured in a three-dimensional system, where cells were seeded as single cells, and colonies analyzed after three weeks (Supplementary Fig. S4A). We found no difference in the number of colonies formed (Supplementary Fig. S4B), but MOXD1 overexpressing cells formed smaller colonies (Supplementary Fig. S4C). This reduction in proliferative capacity was not reflected in cells grown in two-dimensional culture system (Supplementary Fig. S4D). No difference was observed in the rate of active Caspase-3 (Supplementary Fig. S4E). Inducing neuronal differentiation is a common way to target NB, directing cells into a less aggressive state (*39*). There were no differences in differentiation between cells with or without MOXD1 expression, as assessed by quantifying the number of neurites per cell as a proxy (Supplementary Fig. S4F-G). Treatment with cisplatin and doxorubicin, two of the first line chemotherapeutic drugs for NB patients (*40*), showed no differences in therapy resistance (Supplementary Fig. S4H-I), and wound healing experiments did not detect any differences in cell motility (Supplementary Fig. S4J-K). We also investigated angiogenic capacity by co-cultures with mouse MS1 endothelial cell. There was no pronounced difference in master segment length (Supplementary Fig. S4L-M).

### MOXD1 signature genes relate to processes in embryonic and tumor development

To elucidate the transcriptional changes associated with MOXD1 we performed RNAseq of *in vitro* grown cells. We retrieved 94 differentially expressed genes (DEGs) when comparing SK-N-BE(2)c^CTRL^ and SK-N-BE(2)c^MOXD1-OE^ (Supplementary Fig. S5A). The *in vitro* grown cells were, as described above, implanted into mice to form tumors. At experimental endpoint (volume >1800mm^3^), tumors were dissected and subjected to RNAseq. Expression of *MOXD1* was enriched in both data sets, confirming maintained overexpression during *in vivo* growth (Supplementary Fig. S5A-B). We retrieved a list of 117 DEGs generated from the *in vivo* RNAseq (Supplementary Fig. S5B). Gene ontology analysis revealed enrichment in genes connected to embryonic development and several processes involved in tumor growth, including extra-cellular matrix and Notch signaling (Supplementary Fig. S5C).

### Knockout of MOXD1 increases tumor growth and penetrance in a NB zebrafish model

Analysis of recently published expression data from the *TH-MYCN* driven NB mouse model (*41*) showed that *MOXD1* expression steadily decreases with tumor progression (hyperplasia at week 2 and malignant tumor at week 6) (Fig. 5A). Mice without tumors (WT) maintained *MOXD1* expression levels above baseline over time (Fig. 5A).

**Fig. 5.**
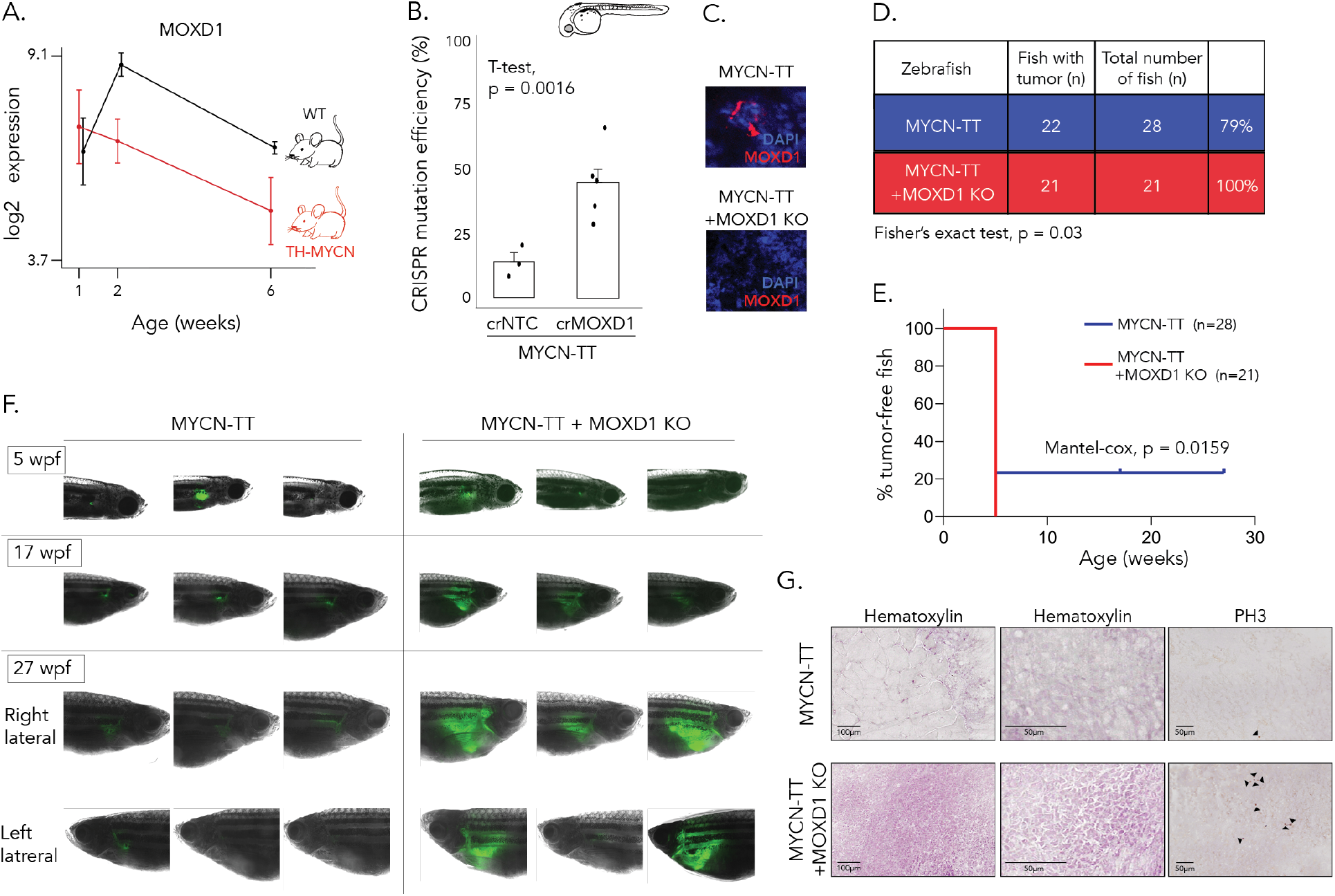
MOXD1 knockout accelerates tumor penetrance in zebrafish. (**A**) *MOXD1* expression score analysis by RNA sequencing (data from De Wyn *et al* (2021)) in wild type (WT) mice without tumors and in TH-MYCN mice (*n*=4 for each group and time point). Graph shows mean expression score ± standard deviation. (**B**) CRISPR mutation efficiency in MYCN-TT+MOXD1 KO and MYCN-TT zebrafish as determined by tumor sample analysis on Miseq and analyzed using CRISPResso2.0 software (http://crispresso2.pinellolab.org/submission). (**C**) Confirmation of CRISPR/Cas9-mediated knockout (KO) of MOXD1 at protein level in tumors dissected from MYCN-TT only and MYCN-TT+MOXD1 KO zebrafish. Staining of MOXD1 by immunofluorescence. DAPI was used to counterstain nuclei. (**D**) Summarizing table of MYCN-TT and MYCN-TT+MOXD1 KO zebrafish with tumors. *p*-value by Fisher’s exact test as indicated. (**E**) Kaplan-Meier plot of tumor-free zebrafish up to experiment endpoint (27 weeks). Number of fish and *p*-value by Mantel-cox as indicated. (**F**) Representative images of zebrafish at 5-, 17- and 27-weeks post-fertilization (wpf). At week 27, each fish has been photographed from both sided (right and left lateral side, respectively). Cancer cells are visualized by fluorescence. Note that images are not from corresponding fish at each time point, but randomly picked from the whole population. (**G**) Hematoxylin staining to visualize tissue of MYCN-TT and MYCN-TT+MOXD1 KO zebrafish tumors as well as staining with phospho-Histone H3 (PH3) visualizing mitosis. Arrowheads denote mitotic cells.

To investigate tumor growth and penetrance in another highly relevant vertebrate *in vivo* system, we utilized the Tg(*dbh:MYCN*; *dbh:EGFP*) zebrafish model, further on described as MYCN-TT (Fig. 5B-G). These fish co-express EGFP and human MYCN under control of the zebrafish *dβh* promoter, and present with a tumor penetrance of 79% (Fig. 5D) (*42*). We knocked out MOXD1 with high CRISPR mutation efficiency (Fig. 5B) and ablated MOXD1 protein levels as determined in tumors dissected 27 weeks post-fertilization (wpf) (Fig. 5C). Notably, CRISPR/Cas-mediated knockout of MOXD1 in the MYCN-TT zebrafish during the embryonic stage increased tumor penetrance to 100% (n=21/21) (Fig. 5D-E). All MYCN-TT + MOXD1 KO fish had developed tumors already after 5 weeks (Fig. 5E). Analyses of the fish at 5-, 17-, and 27-wpf showed a remarkable difference in tumor size (visualized by fluorescence microscopy; Fig. 5F). Sectioning of dissected tumors at 27wpf revealed differences in tissue architecture where MYCN-TT-derived tumors consisted of large amounts of connective tissue and contained smaller foci of cancer cells (Fig. 5G), while MYCN-TT + MOXD1 KO-derived tumors largely consisted of cancer cells (Fig. 5G). The proportion of proliferating cells (as measured by PH3 expression) was higher in MOXD1 KO tumors (Fig. 5G).

### Conditional knockout of MOXD1 disrupts tissue homeostasis and organ formation

Since MOXD1 is endogenously enriched in tNC during development (Fig. 3), we created CRISPR/Cas-mediated conditional knockouts using the chick embryo. Tissue-specific knockout of MOXD1 was achieved by injection into the lumen of the neural tube, targeting tNCCs at stages before delamination and migration (embryonic day E1.5; Fig. 6A). To investigate effects on organogenesis, we harvested embryos at E6, E10 and E15, and sectioned transversally throughout the trunk. Analysis of MOXD1 in adrenal gland (AG) and sympathetic ganglia (SG), homes for NB, showed that expression remained absent in knockout embryos (Fig. 6B). Trunk NCCs start to express HNK1 post-EMT and delamination from the neural tube. At all developmental stages, we find migrating HNK1^+^ cells along sympathetic ganglia (Fig. 6C). During the time as AG is forming at E6, we detect tNCCs in both groups, however, the AG structure is slightly smaller in knockout embryos (Fig. 6D). At E10 and E15, the AG is visible by HNK1 expression and its characteristic triangular shape in control embryos. In knockout embryos however, tNCCs do not reach, but instead migrate alongside, the AG (Fig. 6D). In addition, the AG has lost its structured shape (Fig. 6D). To investigate where the HNK^+^ tNCCs migrate in knockout embryos, we analyzed tissues additional to AG. While in control embryos there were few positive cells in the Organ of Zuckerkandl (ZO), we found a large amount of invading tNCCs in this area in knockout embryos at E10 (Fig. 6E). At E15, control embryos had retained their ZO morphology, while HNK1^+^ cells collected in foci dispersed across ZO (Fig. 6E). Tyrosine hydroxylase (TH) is a marker of neuroblasts (TH^+^) and chromaffin cells (TH^++^) of SG and AG. As expected, these tissues are composed of TH positive cells in control embryos (Fig. 6F-G), but only sparse positive cells in the AG of knockout embryos (Fig. 6F). In addition, SG form also in knockout embryos, but are smaller than their control counterparts (Fig. 6G). As expected, there were no TH+ cells in the ZO (Fig. 6H).

**Fig. 6.**
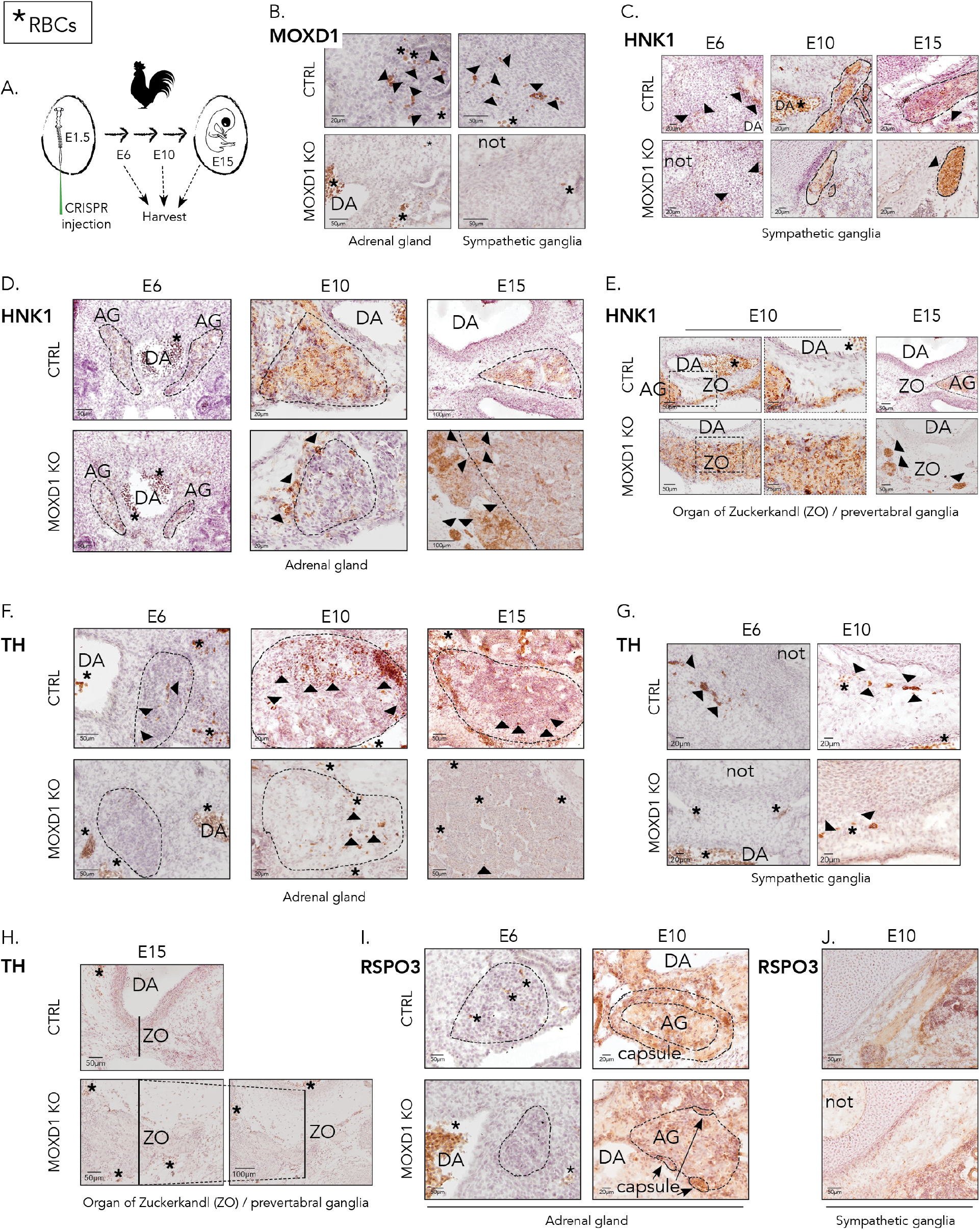
Knockout of *MOXD1* disrupts adrenal gland formation. RBCs (red blood cells; auto-reflecting erythrocytes) are marked by asterisk (**\***). **(A)** Schematic description of the experiment with CRISPR/Cas9-mediated conditional knockout (KO) of *MOXD1* in trunk neural crest cells. Chick embryos were injected with DNA and electroporated by microinjection into the neural tube at E1.5. Embryos were dissected at E6, E10, and E15, sectioned at the trunk axial level, and stained. **(B)** Confirmation of CRISPR/Cas9-mediated KO of MOXD1 at protein level. The trunk part of the chick embryonic body was harvested at E6 (*i.e*., 4.5 days post-injection) and immunohistochemically stained for MOXD1. **(C)** Staining of HNK1 in sympathetic ganglia of chick embryos harvested at E6, E10, and E15, to visualize cells descendent from trunk neural crest. **(D)** Staining of HNK1 in adrenal gland of chick embryos harvested at E6, E10, and E15, to visualize cells descendent from trunk neural crest. **(E)** Staining of HNK1 in organ of Zuckerkandl of chick embryos harvested at E10 and E15, to visualize cells descendent from trunk neural crest. Right images at E10 are magnifications of left images. **(F)** Staining of TH in adrenal gland of chick embryos harvested at E6, E10, and E15, to visualize adrenal gland morphology. **(G)** Staining of TH in sympathetic ganglia of chick embryos harvested at E6 and E10, to visualize sympathetic ganglia morphology. **(H)** Staining of TH in organ of Zuckerkandl of chick embryos harvested at E15, to visualize the absence of the protein in this structure. **(I)** Staining of RSPO3 in adrenal gland of chick embryos harvested at E6 and E10, to visualize the capsule surrounding the adrenal gland. **(J)** Staining of RSPO3 in sympathetic ganglia of chick embryos harvested at E10, to visualize the absence of capsule surrounding this structure. (B-J) Arrowheads indicate cells positive for respective protein. DA, dorsal aorta; not, notochord; AG, adrenal gland; ZO, organ of Zuckerkandl/pre-vertebral ganglia.

### Knockout of MOXD1 disrupts adrenal gland encapsulation in the chick

We further stained embryos for RSPO3, a protein expressed in the capsule of the AG. We did not detect any RSPO3 positive cells in the forming AG at E6 (Fig. 6I). At E10, the AG of control embryos is encapsuled with RSPO3^+^ cells, however, this structure is disrupted in knockout embryos with only fractions of encapsulation (Fig. 6I). To ensure capsule-specificity in the AG, we also stained SG for RSPO3 and could not detect any positive cells (Fig. 6J). To investigate the proliferative rate of these structures, we stained embryo sections for phospho-Histone H3 (PH3) and found small fractions of positive cells in the forming-(E6) and in the developed AG (E10) in control embryos (Fig. 7A). In conjunction with no HNK1^+^ and only few TH^+^ cells in the knockout embryonic AG, there were virtually no proliferating cells in this structure (Fig. 7A). There were only occasional proliferating cells in ZO (Fig. 7B).

**Fig. 7.**
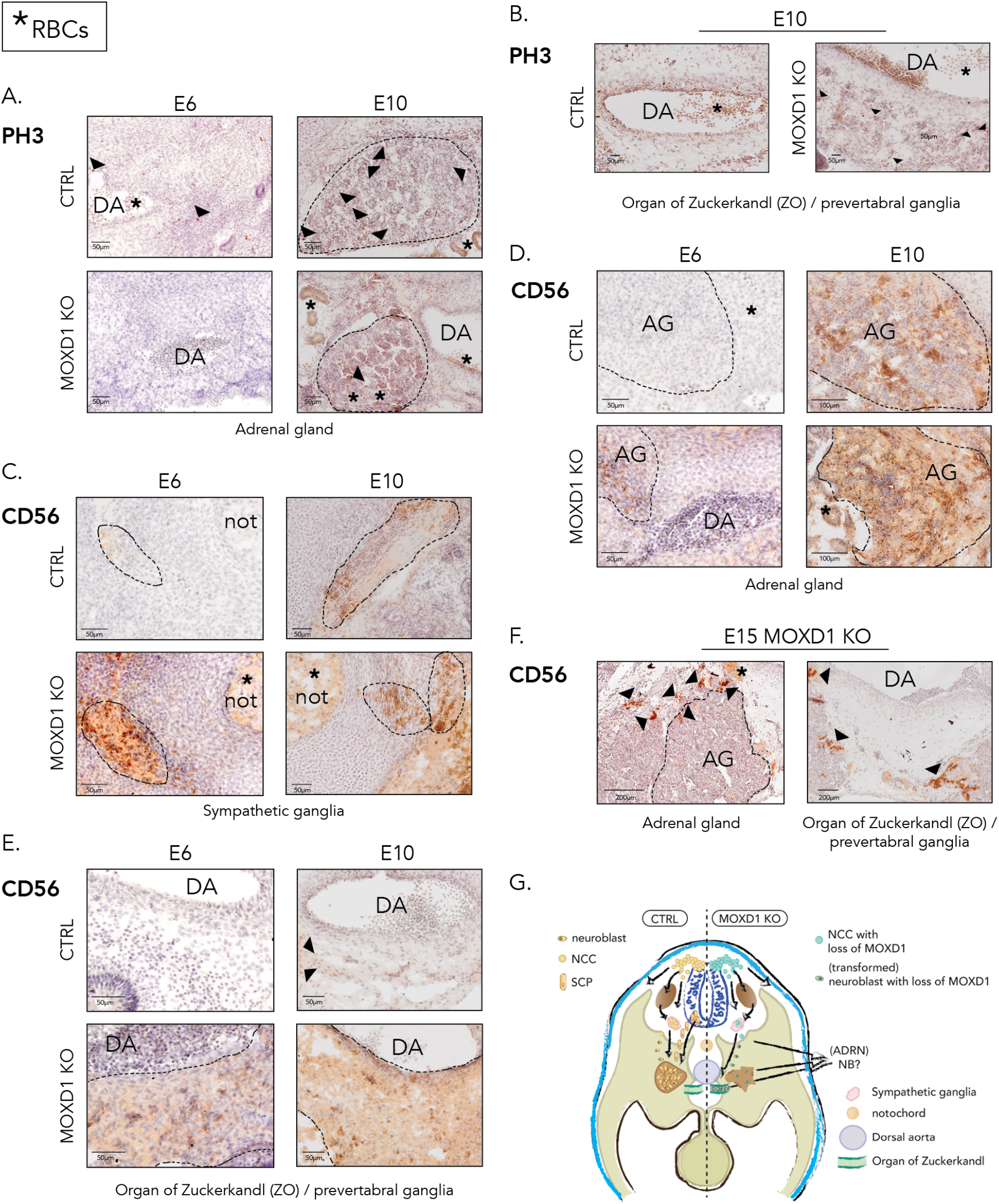
Neural crest cells with MOXD1 knockout express CD56. **(A)** Staining of phospho-Histone 3 (PH3) in adrenal gland of chick embryos harvested at E6 and E10, to visualize mitotic cells. **(B)** Staining of phospho-Histone 3 (PH3) in organ of Zuckerkandl of chick embryos harvested at E10, to visualize mitotic cells. **(C)** Staining of CD56 (NCAM) in adrenal gland of chick embryos harvested at E6 and E10. **(D)** Staining of CD56 in sympathetic ganglia of chick embryos harvested at E6 and E10. **(E)** Staining of CD56 in organ of Zuckerkandl of chick embryos harvested at E6 and E10. **(F)** Schematic summary of the effects on organogenesis from conditional trunk neural crest cell knockout of MOXD1. (A-E) Note that there is background staining from red blood cells (RBCs) in sectioned embryos, marked by asterisk (**\***). Arrowheads indicate cells positive for respective protein. DA, dorsal aorta; not, notochord; AG, adrenal gland; ZO, organ of Zuckerkandl/pre-vertebral ganglia.

### MOXD1 knockout cells express high levels of NB marker CD56

Embryonic and adult AG express the cell adhesion molecule NCAM/CD56 (*43*, *44*), and this protein is clinically used as a diagnostic marker for NB. We found only few and weakly stained CD56^+^ cells in the AG and SG of control embryos at E6 (Fig. 7C-D). At E10, the AG and SG were composed of a substantial fraction of CD56^+^ cells, however these were mixed with cells lacking expression of this molecule (Fig. 7C-D). In contrast, we detected CD56^+^ cells in the AG of embryos with knockout of MOXD1 already at E6 (Fig. 7C). At E10, the AG of knockout embryos were completely filled with CD56^+^ cells, and this was seen also in the SG at both E6 and E10 (Fig. 7C-D). We further analyzed the ZO and while there were virtually no positive cells in control embryos, the majority of cells in MOXD1 knockout embryos expressed CD56 (Fig. 7E). At E15, in coherence with the location of HNK^+^ NC-descendent cells (Fig. 6D-E), MOXD1 knockout results in collection of strongly expressing CD56^+^ cells outside of the AG and in the ZO (Fig. 7F).

In conclusion, the tissue architecture of AG is severely disrupted following conditional MOXD1 knockout in pre-EMT tNCCs, and these cells migrate alongside the AG and unconventionally collect in the ZO (Fig. 7G).

## Discussion

This study establishes *MOXD1* as a *de novo* tumor-suppressor gene in NB. We employ several highly relevant vertebrate model organisms (zebrafish, chick, mouse, and human), bulk RNAseq of representative cohorts of the clinical spectra, scRNAseq of normal adrenal gland and adrenal gland-derived NB, NB-derived cell cultures, as well as tissue material from NB patients, and show translatability of our results across species and thus relevance for human disease. The overlapping results in our developmental models reinforces that the functions of MOXD1 could be conserved along the vertebrate lineage.

Like with most childhood cancer types, few NBs are mutation-driven. In conjunction, we find no mutations in the *MOXD1* gene, however, 4-5% of all NBs display loss of chromosomal 6q23, surrounding the *MOXD1* locus. Loss of MOXD1 in high-risk NB patients with an already dismal prognosis remarkably reduces the survival probability from 40% to under 20%. We show with experimental *in vivo* studies in mice, zebrafish and chick embryos (in the latter exploiting the CAM assay) that overexpression of MOXD1 in NB cells without endogenous expression of the protein delays tumor growth and increases survival. In contrast, our data shows that MOXD1 knockout instead accelerates tumor penetrance, tumor formation and decrease embryo survival.

A recent study of non-tNC-derived tumor form glioblastoma identified that high MOXD1 expression associated with poor prognosis (*45*), which is opposite to our findings in NB. To explore the prognostic role of MOXD1 across cancer types, we therefore analyzed patient data from two additional non-tNC-derived tumor forms: breast and colon. We found that MOXD1 expression does not discriminate between localized and aggressive disease in any of these cancer types. Malignant melanoma is alike NB a tNC-derived cancer, and to understand whether MOXD1 can predict outcome in all tNC-derived cancers, or only NB specifically, we compared *MOXD1* expression in melanoma and NB. We found that MOXD1 expression is high in melanoma as compared to NB and that MOXD1 levels are slightly higher in metastatic as compared to primary melanoma. Although NB and melanoma are both tNC-derived, they descend from cells migrating along different pathways (ventral and dorsal, respectively) following delamination and EMT. Thus, we hypothesize that the role of MOXD1 in cancer is tissue- and cell-specific, and even depend on lineage commitment within the same embryonic progenitor cell population. This is strengthened by the results that MOXD1 is expressed almost exclusively in SCPs during healthy embryogenesis. Two recent studies provide in-depth scRNAseq data from human and mouse neural crest development (*14*, *36*). Although these reports substantially contribute to our understanding of cell commitment and differentiation, they focus on developmental stages after sympathoblasts have left from the neural crest (SCPs from E11.5-E15.5 and adrenal gland post-conception day (PCD) 50-120 in mouse and human, respectively). In fact, NCCs have completed their migration by E10.5 in mice and PCD 40 in humans, and available data do thus not capture events that take place during EMT and migration. Our model of conditional knockouts in chick embryos, initiated at stages before NCCs delaminate and migrate, and with the possibility to capture cells all the way up to hatching, is an advancement in the field that opens up for lineage tracing from the very earliest events. Considering our data showing that MOXD1 plays a role in NB but found no such correlations in melanoma, two tNC-derived cancers, specifically highlight the importance of studying cells before the process of EMT and induced migration.

The two distinct cellular phenotypes that foster NB development and progression are immature MES and lineage-committed ADRN. The MES core signature is highly enriched in SCPs, and our finding that MOXD1 is expressed in MES, but not ADRN, cells point to the fact that MOXD1 is lineage-restricted to SCPs during healthy embryogenesis, and might impact the subtype and aggressiveness of the resulting tumor. At this point, it is however unclear whether MOXD1 directs lineage commitment where loss of MOXD1 skews cells towards a noradrenergic differentiated ADRN phenotype, or MOXD1 is lost as a consequence of ADRN tipping.

In conjunction with the distinct difference in MOXD1 levels between ADRN and MES phenotypic cells, overexpression in cells without, and knockout in cells with endogenous protein expression have a major impact on tumor formation, growth and survival. ADRN-like NB cells drive tumor population whereas MES-like cells are resistant to therapy. The overexpression of MOXD1 in ADRN NB cells might shift cells towards a more MES-like state and thus explain them being less aggressive *in vivo*. It remains debated whether ADRN and MES are distinct cell types or states during the life cycle of a cell, if a potential homogenous MES cell type exists, and whether the mechanisms and frequency of possible transdifferentiation between the two types or states occur *in vivo*. Regardless, these results do not settle where MOXD1 plays a role during the tumorigenic process. We hypothesize that MOXD1 functions as a tumor suppressor gene during priming and/or initiating event, due to the facts that; (1) MOXD1 knockout in zebrafish models of NB increases tumor penetrance to 100%, (2) MOXD1 expression does not affect response to cytostatic drugs (late event), and (3) conditional knockout in healthy progenitor cells affects organogenesis, tissue architecture, and cellular localization.

SCPs generate the majority of chromaffin cells and a smaller portion of neuroblasts composing the adrenal medulla (*15*, *36*). However, a recent study debate whether these findings are true only in younger patients (<18 months at diagnosis). Bedoya *et al* suggest a SCP-derived subtype of NB in younger children, accompanied by better prognosis (*7*). Applying scRNAseq data from the developing adrenal gland from both mouse and human, we could show that *MOXD1* expression is virtually restricted to SCPs during development, excluding expression in cells of later stages: bridge cells, chromaffin cells and sympathoblasts. MOXD1 overexpression-moderated extended survival *in vivo* and reduced overall tumor burden, *i.e*., generating a less aggressive tumor phenotype, is possibly mediated by directing cells towards a more SCP-like phenotype.

The sparse literature on the functions of MOXD1 that exists suggests involvement of binding copper (*46*). Disrupted copper homeostasis is known to be involved in malignant progression of several tumor forms (*47*, *48*). Both increased and decreased levels of copper-related proteins have been connected to cancer and their use as predictive or prognostic biomarker is promising (*49*). However, additional studies with further functional and biological experiments are needed to fully unravel the mechanisms of MOXD1. In conclusion, we identify *MOXD1* as a novel tumor suppressor gene and lineage-specific marker for immature cells during embryogenesis as well as in NB. Our results suggest that NB subtypes exist and derive from restricted cell lineages during healthy neural crest development.

## Acknowledgments

The authors would like to thank Christina Möller for excellent technical assistance. The authors would like to acknowledge Clinical Genomics Lund, SciLifeLab and Center for Translational Genomics (CTG), Lund University, for providing expertise and service with sequencing and analysis. The authors would like to thank the Zebrafish Facility Ghent (ZFG) Core at Ghent University, Belgium.

## Funding

Swedish Cancer Society (SM, EHa, MAH)

The Swedish Childhood Cancer Fund (SM, MAH)

The Crafoord Foundation (SM, MAH, EHa)

The Jeansson Foundations (SM)

The Ollie and Elof Ericsson Foundation (SM, EHa)

The Magnus Bergvall Foundation (SM)

The Hans von Kantzow Foundation (SM)

The Royal Physiographic Society of Lund (SM, GAH, SK, TG)

The Gyllenstierna Krapperup’s Foundation (SM)

The Franke and Margareta Bergqvist Foundation (SM)

Gunnar Nilsson Cancer Foundation (SM)

Mrs Berta von Kamprad Foundation (SM)

Wenner-Gren Foundations (GAH)

FWO Vlaanderen (Fund for scientific research Flanders) (EHi)

Innovative Therapies for Children with Cancer (SL)

Villa Joep (FS)

Kinderkankerfonds (FS)

Fonds Oudaan vzw (FS)

Geconcenteerde onderzoeksactie (GOA), Ghent University: Replication fork protector dependency factors as novel targets for combination treatment and immunomodulation in neuroblastoma (FS)

FWO research project (FS)

## Author contributions

Conceptualization: EF, SA, SM

Methodology: EF, SA, EHi, GAH, EM, SL, JWB, EMi, RN, FS, JvN, and SM

Validation: EF, SA, EHi, GAH, SM

Formal Analysis: EF, SA, EHi, GAH, EMo, JvN, SM

Investigation: EF, SA, EHi, GAH, EMo, SK, TG, SL, JWB, EL, EMa, EMi, MC, MK, NE, JvN, SM

Resources: RN

Data Curation: SA, JvN, SM

Visualization: EF, SA, EHi, GAH, JvN, SM

Funding acquisition: SM

Project administration: SM

Supervision: FS, RN, SM

Writing – original draft: EF, SA, EHi, GAH, EMo, SM

Writing – review & editing: SK, TG, SL, JWB, EL, EMa, EMi, MC, MK, NE, CM, MAH, EHa, RN, FS, JvN, SM

## Data and materials availability

Sequencing data will be made available upon acceptance.

## Supplementary Materials

Materials and Methods

Figs. S1 to S5

Tables S1 to S2

