## Supplementary Materials, Figures and Tables for "MOXD1 is a gate-keeper of organ homeostasis and functions as a tumor-suppressor in neuroblastoma"

**This PDF file includes:**

Materials and Methods
Figs. S1 to S5
Tables S1 to S2

### Materials and Methods

#### *Ethics*

According to Swedish regulation (Jordbruksverkets föreskrift L150, §5), experiments performed on chick embryos older than embryonic day 13 require ethical permit. All chick embryo and mouse procedures followed the guidelines set by the Malmö-Lund Ethics Committee for the use of laboratory animals and were conducted in accordance with European Union directive on the subject of animal rights (ethical permit nr. 18743/19). For the TMA, all patients, their relatives or their legal guardians signed the appropriate written informed consent. The present study was approved by INCLIVA's Clinical Research Ethics Committee (ref. 2017/198). All procedures involving zebrafish were approved by the local animal ethics committee (Ghent University hospital, Ghent, Belgium; ethical permit nr. ECD 17/100) and performed according to local guidelines and policies in compliance with national and European law. Human embryonic tissue was obtained from elective abortions under ethical permit Dnr 6.1.8-2887/2017, Lund University, Sweden.

#### *Cell culture*

The neuroblastoma cell lines SK-N-BE(2)c, SK-N-SH, SH-SY5Y, SH-EP, and IMR-32 (kind gifts from Drs. June Biedler, Memorial Sloan Kettering Cancer Institute and Robert Ross, Fordham Unibversity) were cultured in MEM or RPMI 1640 (IMR-32), supplemented with 10% fetal bovine serum. 691-ADRN were cultured in DMEM/F12 supplemented with 40 ng/ml FGF, 20 ng/ml EGF, 1x B27 supplement. MS1 mouse endothelial cells were cultured in DMEM supplemented with 10% fetal bovine serum. Penicillin (100 units) and streptomycin (10 µg/mL) were added to all cultures. Cells were kept in humidified incubator at 37°C, at 21% $O_2$, and 5% $CO_2$. SK-N-BE(2)c, SK-N-SH, SH-SY5Y, SH-EP, IMR-32, and MS1 were grown as monolayer and dissociated using trypsin and accutase. 691-ADRN were grown in suspension and dissociated using trypsin. All cells were at minimum tri-monthly screened for mycoplasma (Eurofins Genomics). The 691 isogenic pair (2018), and SK-N-BE(2)c, IMR-32, SH-SY5Y, and SH-EP cell lines (all 2022) were authenticated by STR profiling (Eurofins Genomics).

#### *Lentiviral transduction*

SK-N-BE(2)c, SK-N-SH and ADRN-691 cells were transduced with lentiviral vector encoding MOXD1 gene or control plasmid (OriGene, RC205862L2; PS100071). Lentiviruses were directly added to the cell culture media and incubated for 16 hours before media change.

#### *Colony formation in soft agar*

SK-N-BE(2)c cells were plated at $2.5 \times 10^3$ cells per well in a mix of DMEM and 0.3 % agar on top of a coating layer of 0.6 % agar in 6-well tissue culture plates. Plates were incubated at 37°C in humidified atmosphere for 21 days. Colonies were stained with crystal violet (0.001 %; Sigma-Aldrich, #C0775). Colony number and size were automatically calculated with ImageJ. Four biologically independent replicates were used for quantification and statistical analysis.

#### *Cell viability and drug response assay*

SK-N-BE(2)c cells were plated in technical triplicates in opaque 96-well plates in a total volume of 100 µL, and thereafter incubated for 24 hours before treatment to allow attachment to the plate. Cells were treated with the indicated logarithmic concentration ranges of cisplatin (Sigma-Aldrich,

#C2210000) and doxorubicin (Sigma-Aldrich, #D1515) dissolved in sterile $H_2O$. Cell viability was measured using CellTiter-Glo (Promega Corp., #G7571). Three biologically independent replicates were used for quantification and statistical analysis.

***Cell proliferation and differentiation***

SK-N-BE(2)c cells were seeded at $1\times 10^5$ cells per well in a 6-well plate, incubated for 5 days, and manually counted using a Bürker chamber. Four biologically independent replicates were used for quantification. For differentiation assay, cells were grown for 24 hours and number of neurites were manually assessed by blinded quantification from representative images. The number of neurites were normalized to total number of cells (*i.e.,* neurites per cell). Three biologically independent replicates were used for quantification.

***Wound healing***

Cells were seeded at $3 \times 10^5$ cells per well in a 6 well plate and incubated at 37°C for 24 hours (for the cells to reach 80% confluency) before a scratch or "wound" was made in the middle of the culture with a p1250 pipette tip. Images were captured at 0, 24, 48, and 72 hours after the scratch. Four biologically independent replicates were used for quantification. Open areas were manually marked in ImageJ and then automatically measured.

***In vivo tumorigenicity***

Female athymic mice (NMRI-Nu/Nu strain; Taconic) were housed in a controlled environment. A total of $0,8 \times 10^6$ SK-N-BE(2)c, $1 \times 10^6$ SK-N-SH, or $0,7 \times 10^6$ 691-ADRN cells were subcutaneously injected in a 100 µL 2:1 mixture of cell culture media and growth factor reduced Matrigel (BD, 354230) to the right flank of mice. Mice were euthanized when tumors reached a size of >1800 $mm^3$ in diameter or 12 months after experiment start (endpoint, 365 days). Tumor pieces were snap-frozen and further processed for RNA sequencing.

***RNA sequencing***

RNA was extracted from harvested cells in culture (n=3 biologically independent samples per group) or pieces from *in vivo* grown tumors (n=5 biologically independent samples per group). Sequencing was performed using NovaSeq 6000 (Illumina). Alignment of reads was performed using the HISAT2 software and the reference genome was from the Ensemble database (Human GRCh38, GTF 94 (*in vitro*) or 99 (*in vivo*)). Expression counts were performed using the StringTie software and differentially expressed gene (DEG) analysis was performed using DESeq2.

DEGs between the MOXD1 overexpressing SK-N-BE(2)c cells and the control group were identified by a log2foldchange >1.5 and a higher expression in all samples in one group compared to all the samples in the other group (*in vitro*) and log2foldchange >1.5 and a higher expression in >3 samples in one group compared to all the samples in the other group (*in vivo*). Gene ontology (GO) enrichment analysis of the DEGs *in vivo* was performed with the R package clusterProfiler. The organism was set to human, the p-value cut-off was 0.05 and the gene count cut-off was 5.

***Co-culture angiogenesis assay***

MS1 (pancreatic islet mouse endothelial) cells ($10^4$) were mixed with SK-N-BE(2)c cells (1:1 ratio) and seeded in a 96-well plate on top of a layer of growth factor reduced Matrigel (BD Biosciences, #354230) in 1:2 MEM and DMEM medium (1:1). Matrigel was thawed at 4°C overnight, added to 96-well plates, and transferred to 37°C to allow for polymerization prior to seeding. Cells were incubated for 8 hours before four images of each group were captured. Images were analyzed in ImageJ using the Angiogenesis Analyzer plug-in.

***Tissue microarray (TMA)***
50 high risk NB samples with *MYCN* amplification were analyzed in a TMA, consisting of one 1 mm cylinder from each of 50 high risk NBs with *MYCN* amplification. Patient samples included in the TMA were referred to the Spanish Reference Centre for Neuroblastoma Biological and Pathological studies (Department of Pathology, University of Valencia-INCLIVA) between 1999 and 2017. Patients' data were collected by the pediatric oncologists in charge at the hospital of origin and by the clinicians of the Reference Centre for NB clinical studies, and were classified according to the INRG pre-treatment stratification criteria.

Paraffin-embedded TMAs were sectioned (3 µm) and stained for MOXD1 using a DAKO Cytomation Autostainer Plus with the EnVision FLEX High pH kit (Agilent, code K8010). Slides were dried at 60°C for 1 hour, paraffin was removed, and slides were placed in a pressure cooker for 20 minutes at 97°C in the EnVision FLEX Target Retrieval Solution Low pH (pH 6; Agilent, code K8005). Slides were washed in Wash buffer (Agilent, code K8007), after which EnVision FLEX Peroxidase Blocking Reagent (Agilent, code DM821) was added. Slides were incubated for 5 minutes and then washed as above. Primary antibody was diluted in EnVision FLEX Antibody Diluent (Agilent, code K8006) and slides were incubated for 30 minutes after washing as above. Slides were then incubated in EnVision FLEX/HRP (Agilent, code DM822), washed as above, and further incubated in EnVision FLEX DAB+Chromogen (Agilent, code DM827) diluted in EnVision FLEX Substrate Buffer (Agilent, code DM823). After a last washing step, slides were incubated with HTX for 3 minutes, dehydrated, and mounted.

Sections were digitalized with the PannoramicMIDI 3DHistech scanner at 20X magnification. Cytoplasmic brown staining was considered MOXD1 positive result. The number of MOXD1 positive cells and the staining intensity was evaluated with NuclearQuant module of Pannoramic Viewer software (3DHistech). This module could be applied to non-nuclear stained compartments by adjusting nucleus size settings to detect entire cells when required (nuclear radius 3-15µm) and adjusting color deconvolution settings. Detected artefacts and folded and/or broken regions were considered uninformative and were excluded from the image analysis. Automatically obtained results were validated with optical microscopy by a pathologist and two expert researchers in morphology. The percentage of positive cells in each sample for every MOXD1 intensity expression (low, medium and high) was calculated. For statistical purposes, we defined samples as low (≤ percentile 25) or high (> percentile 25) MOXD1 expression, according to the percentage of positive cells that were digitally detected as presenting high intensity MOXD1 expression. Additionally, we differentiated between homogeneous and heterogeneous MOXD1 samples, considering a sample heterogeneous when its total MOXD1 positive cell population was characterized by having cells with low, medium and high intensity MOXD1 expressions, but none of the positive cell subpopulations represented more than 66% of the total MOXD1 positive cell population. SPSS version 26 was used to perform the statistical analysis, setting the significance level at 95%. Using the Chi-square test we evaluated statistical correlation between MOXD1 expression patterns and patient age (< 18 versus ≥ 18 months). Other INRG prognostic variables could not be tested due to the homogeneous characteristics of the studied cohort.

***Human embryonic tissue***

Human embryos obtained from elective abortions were collected in custom-made hibernation medium (Life Technologies). Embryonic/fetal stages were determined by ultrasound. Embryos were dissected for selected tissue (trunk part of the embryonic body), transferred to 4% PFA and incubated in 4°C overnight. Next day, samples were processed through a 5% to 15% sucrose gradient, incubated in gelatin overnight, and finally embedded and snap-frozen. Embryos were transversally cryosectioned at 7-12μm, de-gelatinized in 42°C for 45-60 min before staining procedures.

***Chick embryos***

Chick embryos were acquired from commercially purchased fertilized eggs and incubated at 37.5°C until desired developmental Hamburger Hamilton (HH) stages were reached[50]. One-sided electroporation *in ovo* were performed with 5 pulses of 30ms each at 22V. Ringer's balanced salt solution containing 1% penicillin/streptomycin was used in all experiments. CRISPR constructs targeting MOXD1 or a gRNA non-targeting control (#99140, Addgene) were electroporated at a concentration of 1.5 μg/μl, accompanied by as Cas9-GFP vector (#99138, Addgene) at 2 μg/μl. All constructs were injected at HH stage 10+/11 into the lumen of the neural tube from the posterior end and embryos were electroporated *in ovo*. Embryos were allowed to sit at room temperature for 6 – 10 hours in order to allow the Cas9 protein to fold. Embryos were then incubated at 37.5°C until desired end points were reached. Embryos were dissected (cranium was dislocated directly after extraction from the egg according to ethical permit, and discarded), fixed in 4% PFA overnight at 4°C, processed through a 5% to 15% sucrose gradient, incubated in gelatin overnight and finally embedded and snap-frozen. Embryos were transversally cryosectioned at 7-12μm, de-gelatinized in 42°C for 45 – 60 min before staining procedures.

***Cloning***

For CRISPR/Cas9 targeting, oligos designed to target *MOXD1* (MOXD1.4.gRNA Top oligo – 5' ggatgGCACCATGTGACAAAGgtg 3', Bot oligo – 5' aaaccacCTTTGTCACATGGTGCc 3') were annealed pairwise at a concentration of 100 μM per oligo using T4 DNA Ligase Buffer by heating to 95°C for 5 minutes. The annealed oligo reactions were cooled to room temperature and diluted 1:1000. The U6.3>gRNA.f+e (#99139, Addgene) vector was digested over night with BsaI-HF enzyme (New England Biolabs, #R3733) and gel extracted. gRNAs were cloned into the digested U6.3>gRNA.f+e vector using T4 DNA Ligase (New England Biolabs, #M0202) at room temperature for 20 minutes. Successful inserts were identified by colony PCR using U6 sequencing primer and gRNA reverse oligo specific to *MOXD1* gRNA.

***Immunohistochemistry***

Human embryos were stained for MOXD1 and TH as described for the TMA. For chick embryos, immunohistochemistry was carried out on transversal cryosections of trunk neural crest. Slides were washed in 0.05% Tween in phosphate saline buffer (PBS-T) for 10 minutes, and incubated with 1ml of hydrogen peroxide (Fischer Scientific, #H/1750/15) in 200 ml of PBS-T to inactivate the endogenous peroxidase activity. Slides were washed in PBS-T two times for 10 minutes each followed by incubation with blocking solution (5% goat serum albumin and 0.5% Triton-X) in a humidified chamber during one hour. Slides were thereafter incubated with primary antibody in a humidified chamber overnight. All primary and secondary antibodies used in the present study are summarize in Supplementary Table S1. Slides were washed three times for 30 minutes each and then incubated with the secondary antibody in a humidified chamber overnight. The samples were washed three times for 10 minutes each and then incubated with streptavidin peroxidase (1:200; #SA-5004, Vector Laboratories) diluted in PBS in a humidified chamber for 2 hours. Finally, the slides were incubated with 0.03% DAB (3,3´diaminobenzidine tetrahydrochloride hydrate, 97%) chromogen and 40µL of hydrogen peroxide in PBS. The reaction was stopped in PBS.

Immunostained slides from chick embryos were washed several times in PBS. Samples from E6 were immersed in hematoxylin colorant (#MHS32, Sigma-Aldrich) for 2 minutes whereas E10 and E15 samples were submerged for 10 minutes followed by washing in PBS. Dehydration in increasing percentages of ethanol was performed followed by exposition to xylene two times for 5 minutes each. Slides were mounted using Eukitt mounting medium (#03989, Sigma-Aldrich).

***RNA extraction and quantitative real-time PCR***

Total RNA from cells was extracted using RNAqueous Micro Kit (Ambion RNAqueous™-Micro Kit, ThermoFisher, #AM1931) and eluted in 20 µL elution solution. cDNA synthesis was performed using random primers (Applied Biosystems Reverse Transcriptase Kit, ThermoFisher, #4368814). Quantitative RT-PCR was performed using SYBR™ Green PCR Master Mix (ThermoFisher, #4364346). Relative mRNA expression was normalized to the expression of three reference genes (*UBC*, *YWHAZ*, and *SDHA*) using the comparative $C_t$ method[51]. Primers are listed in Supplementary Table S2.

***Zebrafish models***

Zebrafish were maintained in a Zebtec semi-closed recirculation housing system (Techniplast, Italy). The water had a constant temperature (27-28 °C), pH (~7.5), conductivity (~500 µS) and light/dark cycle (14h/10h). The fish were fed twice a day with dry food (Gemma Micro, UK) and once with Artemia (Ocean Nutrition, Belgium).

*Determining the best crRNA for MOXD1-KO*

CRISPR RNAs (crRNAs) targeting MOXD1 were designed and injected according to a previously published workflow[52]. First, the online Benchling tool (https://www.benchling.com/crispr) was used to select five MOXD1 crRNAs (CATGTTGGGATGTCAATAGG, CCAGGCATGACGGATTACAT, GATGCTGGAGTCATCGAGAC, GAGGCGCTACGATGCTGGAG, CGCCAAGTGCGAGAGTTTAC) with the highest on-target score (efficiency score) and lowest off-target score (specificity score). These crRNAs (200 µM) were mixed together with a general tracrRNA (200 µM) (IDT #1072533) to generate five different gRNA duplexes. Subsequently, 900 pg gRNA duplex was combined with 900 pg Cas9 enzyme (IDT #1081061) before injection (1.4 nl). A non-targeting crRNA (GCAGGCAAAGAATCCCTGCC) was used as a control. Injections of all five crRNA:tracrRNA-Cas9 mixtures were performed in the yolk of wild-type zebrafish embryos during the one-cell stage. For each crRNA injection mix, DNA was isolated from a pool of 24-48 hour post-fertilization (hpf) embryos and amplified using specific primers surrounding the targeted region. The amplified pooled DNA sample was run on a Miseq and analysed via CRISPResso2.0 to determine the mutation efficiency. crMOXD1_3 (GATGCTGGAGTCATCGAGAC) resulted in the highest efficiency score and was used in subsequent experiments.

*Injection of crMOXD1 in MYCN-TT (Tg(dβh:eGFP;dβh:MYCN)) zebrafish line*

The CRISPR-Cas9 injection mixes of crRNA_MOXD1_3 and negative control were injected in MYCN-TT one-cell stage zygotes. This transgenic zebrafish line was previously established by co-injection of eGFP and human MYCN, both under control of the zebrafish DBH promotor, which allows the selection of MYCN-TT-positive larvae at five days post fertilization (dpf) and the detection of NB formation by eGFP expression in the inter-renal gland of the zebrafish. From 5 wpf, the zebrafish were screened for tumor formation biweekly using fluorescence microscopy (Nikon, SMZ18).

The mutation efficiency of the injections was checked on Miseq in a separate embryo pool (24-48 hpf) and the experiments were only continued when mutation efficiencies were higher than 60%. At 9 wpf, genotyping was performed by fin-clipping to confirm the MYCN-TT-positive genotype. Only MYCN-TT-positive zebrafish were included in the study.

Tumors were harvested at 27wpf (n = 6 crMOXD1, n = 3 crNTC), fixed in Modified Davidson's Fixative (22% Formaldehyde (37%), 12% Glacial acetic acid, 33% Ethanol (95%), 34% distilled water) over night on shaker at room temperature, and stored at -80°C until embedding. Tumor pieces were then directly embedded in gelatin, snap-frozen and sectioned at 8µm before degelatinized and processed for IHC and IF as described above.

***Analysis of publicly available datasets***

The neuroblastoma patient datasets SEQC (GEO:GSE62564), Kocak (GEO:GSE45547), Depuydt (GEO:GSE103123), were downloaded from the R2: genomics analysis and visualization platform (http://r2.amc.nl). The NC/MES/ADRN dataset (R2 ID: ps_avgpres_gsenatgen2017geo52_u133p2), the melanoma dataset (GEO:GSE65904) and the Cancer Cell Line Encyclopedia (GEO:GSE36133) were downloaded from the R2 platform and the NB/GN dataset (GEO:GSE7529), ES/tNC (GEO:GSE109267) breast- (GEO:GSE102484) and colorectal cancer (GEO:GSE39582) were downloaded from GEO. A limma test was conducted on the R2 platform to identify the DEGs between the tumors of the patients belonging to the first and forth quartile of *MOXD1* mRNA expression in the SEQC cohort (*p*-value cut-off 0.05). Survival analysis was performed by log-rank tests with the Kaplan-Meier method and multivariate Cox regressions using the R packages Survival and Survminer (Figs. 1A-C and 1I and Supplementary Fig. 2B). Supplementary Figs. 2C-F were generated using https://padpuydt.shinyapps.io/check_cn_in_hr_nb/, maximum size of aberration was set to 180 Mb. Further analysis of MOXD1 expression in the datasets was conducted in RStudio.

The MOXD1 mRNA expression in the single nuclei data by Bedoya-Reina *et al* (2021) and in the single cell data from Kastriti *et al* (2022) was analyzed with Pagoda at http://oxygen.mtc.ki.se/nc_nb_2021/neuroblastoma/index.html and https://adameykolab.srv.meduniwien.ac.at/glia_gene_umap/.

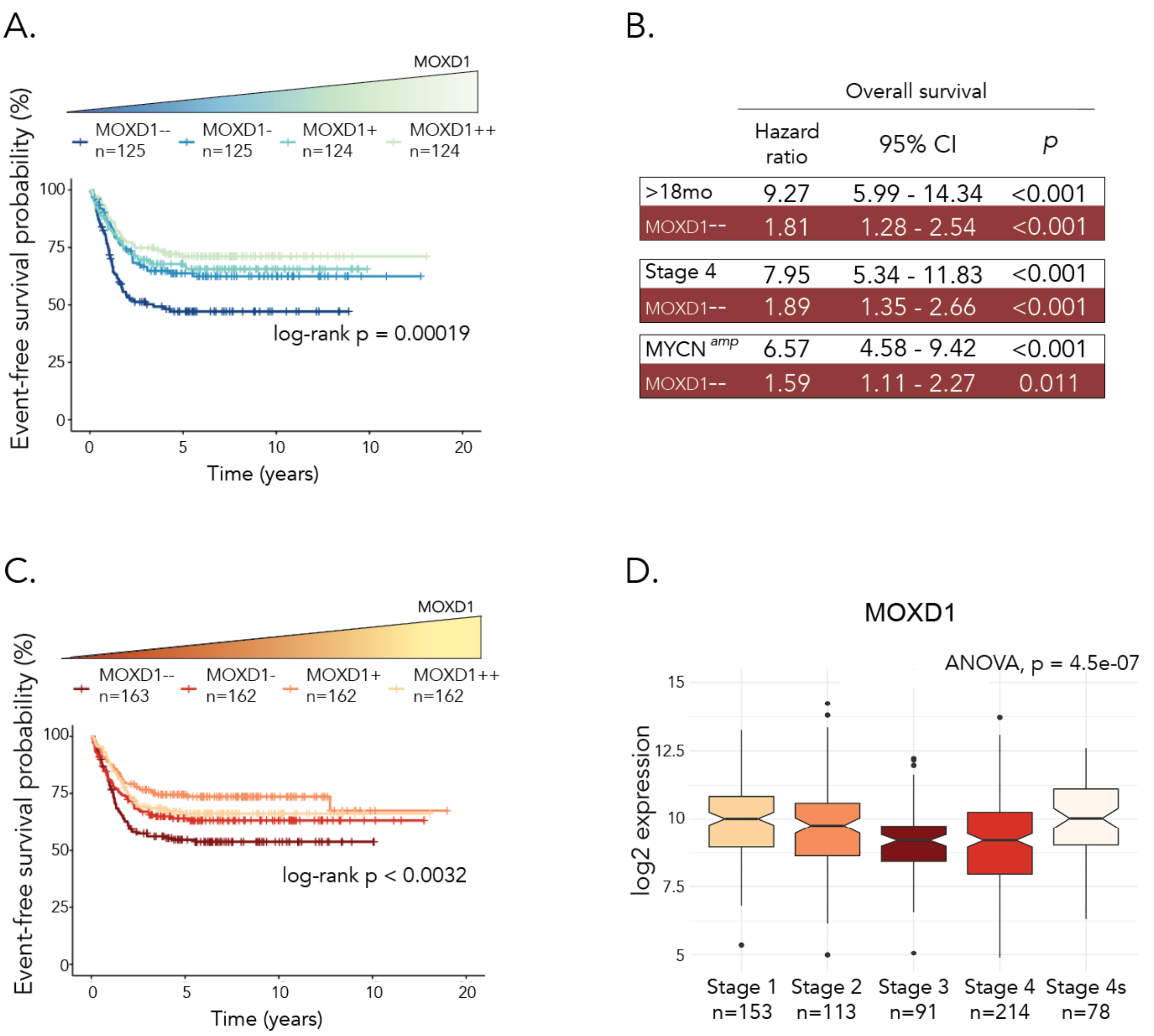

**Fig. S1. Low expression of *MOXD1* correlates to poor outcome in NB.**
**(A)** Kaplan Meier plot of event free survival of NB patients from SEQC cohort ($n = 498$) stratified according to quartiles of *MOXD1* expression. Number of patients ($n$) is specified for each subgroup. $p$-value by log-rank test as indicated. **(B)** Prognostic effect of *MOXD1* mRNA expression for NB patients (Kocak cohort, comparing first and fourth quartile of *MOXD1* expression, see Fig. 1C) with adjustment for age at diagnosis, INSS stage of disease and *MYCN* amplification status, respectively. Hazard ratios, 95% confidence intervals (CI) and log rank $p$-values are given by multivariate Cox regression analyses. **(C)** Kaplan Meier plot of event free survival of NB patients from Kocak cohort ($n$ = 649). Number of patients ($n$) is specified for each subgroup. $p$-value by log-rank test as indicated. (**D**) *MOXD1* expression across the INSS stages in NB patients from Kocak cohort. Number of patients ($n$) is specified for each stage and $p$-value by ANOVA test as indicated.

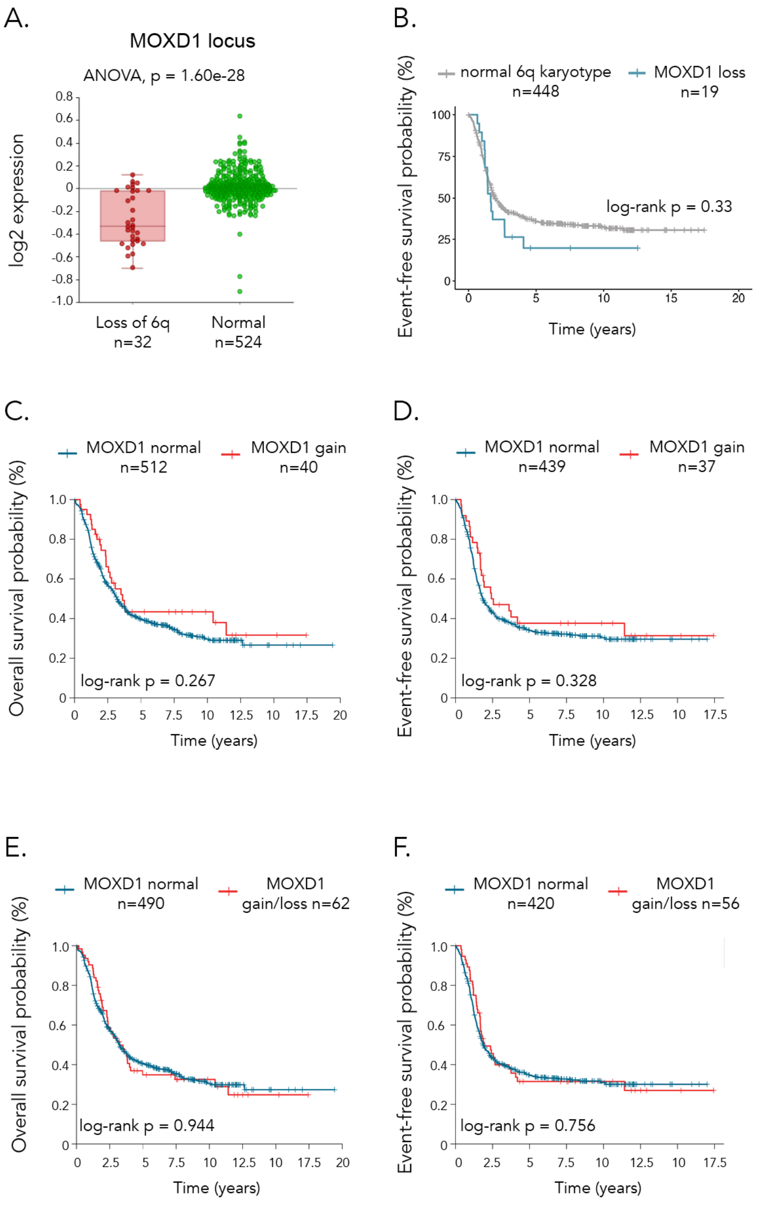

**Fig. S2. MOXD1 locus is lost in high-risk neuroblastoma.**

**(A)** Expression of genes in the surrounding *MOXD1* locus (ARG1, CTGF, ENPP1, ENPP3, STX7, MED23, MOXD1, OR2A4, and CTAGE9) in patients with a normal 6q karyotype as compared to patients with loss of 6q (as defined in, and data, from Depuydt et al (2018)). **(B)** Kaplan-Meier plot of event-free survival of high-risk NB patients (Depuydt; $n$ = 556). Patients stratified by loss of *MOXD1* or no loss of *MOXD1*. Patients with distal 6q mutations not affecting *MOXD1* and patients lacking event-free survival information were excluded. $p$-value by log-rank test as indicated. **(C)** Kaplan-Meier plot of overall survival of high-risk NB patients (Depuydt; $n$ = 556). Patients stratified by gain of *MOXD1* or no loss of 6q (normal). **(D)** Kaplan-Meier plot of event-free survival of high-risk NB patients (Depuydt; $n$ = 556). Patients stratified by gain of *MOXD1* or no loss of 6q (normal). **(E)** Kaplan-Meier plot of overall survival of high-risk NB patients (Depuydt; $n$ = 556). Patients stratified by loss or gain of *MOXD1* or no loss of 6q (normal). **(F)** Kaplan-Meier plot of event-free survival of high-risk NB patients (Depuydt; $n$ = 541). Patients stratified by gain of *MOXD1* or no loss of 6q (normal).

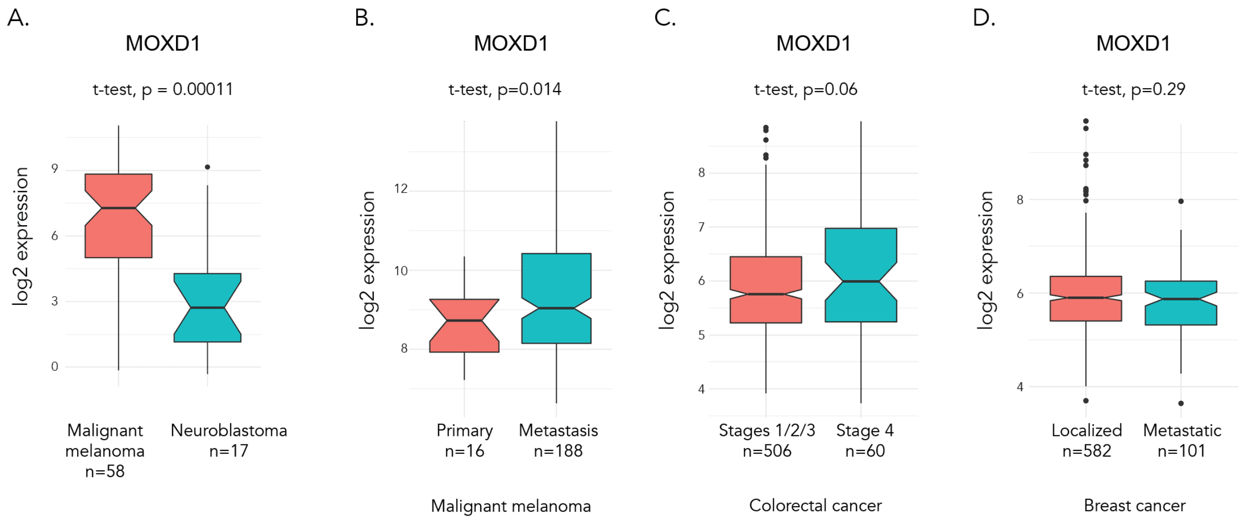

**Fig. S3. MOXD1 as a predictor of poor prognosis is tumor form-specific.**
**(A)** Expression of *MOXD1* in malignant melanoma vs NB, data extracted from the Cancer Cell Line Encyclopedia set. **(B)** Expression of *MOXD1* in primary or metastatic malignant melanoma. Number of patients and *p*-value by t-test as indicated. **(C)** Expression of *MOXD1* in colorectal cancer of Stage 4 vs Stages 1/2/3. Number of patients and *p*-value by t-test as indicated. **(D)** Expression of *MOXD1* in localized vs metastatic breast cancer. Number of patients and *p*-value by *t*-test as indicated.

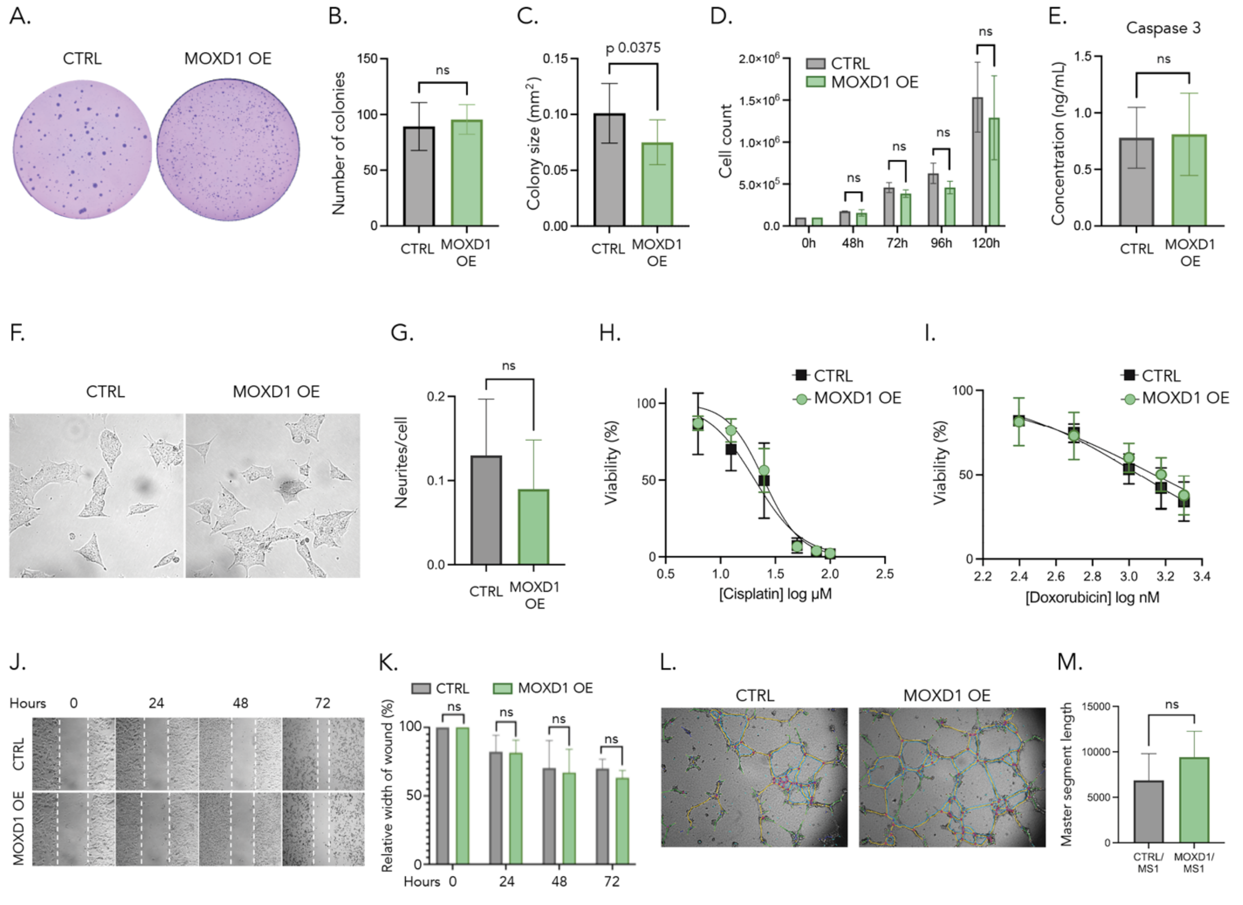

**Fig. S4. Overexpression of MOXD1 does not affect NB cell differentiation or drug response *in vitro*.** (**A**) Representative images from sphere-forming assay of control (CTRL) and MOXD1 overexpressing (OE) SK-N-BE(2)c cells. (**B-C**) Number (B) and size (C) of colonies formed by SK-N-BE(2)c CTRL and MOXD1 OE cells ($n$ = 3 biologically independent repeats for each group). Values are presented as mean and bars indicate standard deviation. $p$-value by unpaired t-test as indicated. ns, $p > 0.05$. (**D**) Proliferative capacity of SK-N-BE(2)c CTRL and MOXD1 OE cells measured by cell count at indicated time points. Values are presented as mean ($n$ = 3 biologically independent repeats for each group) and bars indicate standard deviation. $p$-value by two-way ANOVA as indicated. ns, $p > 0.05$. (**E**) Detection of active caspase 3 as determined by ELISA assay in SK-N-BE(2)c CTRL and MOXD1 OE cells. Concentrations are presented as mean ($n$ = 3 biologically independent repeats for each group and time point) and bars indicate standard deviation. ns, $p > 0.05$. (**F**) Representative brightfield images of SK-N-BE(2)c CTRL and MOXD1

OE cells 24 hours post seeding, quantification in (G). (**G**) Blinded quantification of neurite outgrowth per cell. Values are reported as mean (*n* = 4 biologically independent repeats for each group), bars represent standard deviation. ns, $p > 0.05$. (**H-I**) Dose-response curves of indicated concentrations of cisplatin and doxorubicin in SK-N-BE(2) CTRL and MOXD1 OE cells. Cell viability was determined by CellTiter-Glo 48 hours post-treatment. Values are reported as mean ± SEM, (*n* = 3 biologically independent repeats for each group). No measured point reached $p < 0.05$. (**J**) Wound healing assay of confluent SK-N-BE(2)c CTRL and MOXD1 OE cells. Representative phase contrast images were captured at indicated time points. **(K)** Width of the wound is presented as percentage at each analyzed time point. Bars indicate standard deviation, *n* = 4 biologically independent repeats for each group. Statistical analysis was performed with two-way ANOVA. ns, $p > 0.05$. (**L**) Representative phase contrast images of SK-N-BE(2)c cells co-cultured with MS1 cells at T=8 hours. Color overlay by ImageJ Angiogenesis Analyzer plug-in. **(M)** Images were analyzed for cellular networks using the ImageJ Angiogenesis Analyzer plug-in. Quantification of master segment length, values are presented as mean and bars indicate standard deviation (*n* = 2 biologically independent repeats for each group). Statistical analysis was performed with unpaired t-test. ns, $p > 0.05$.

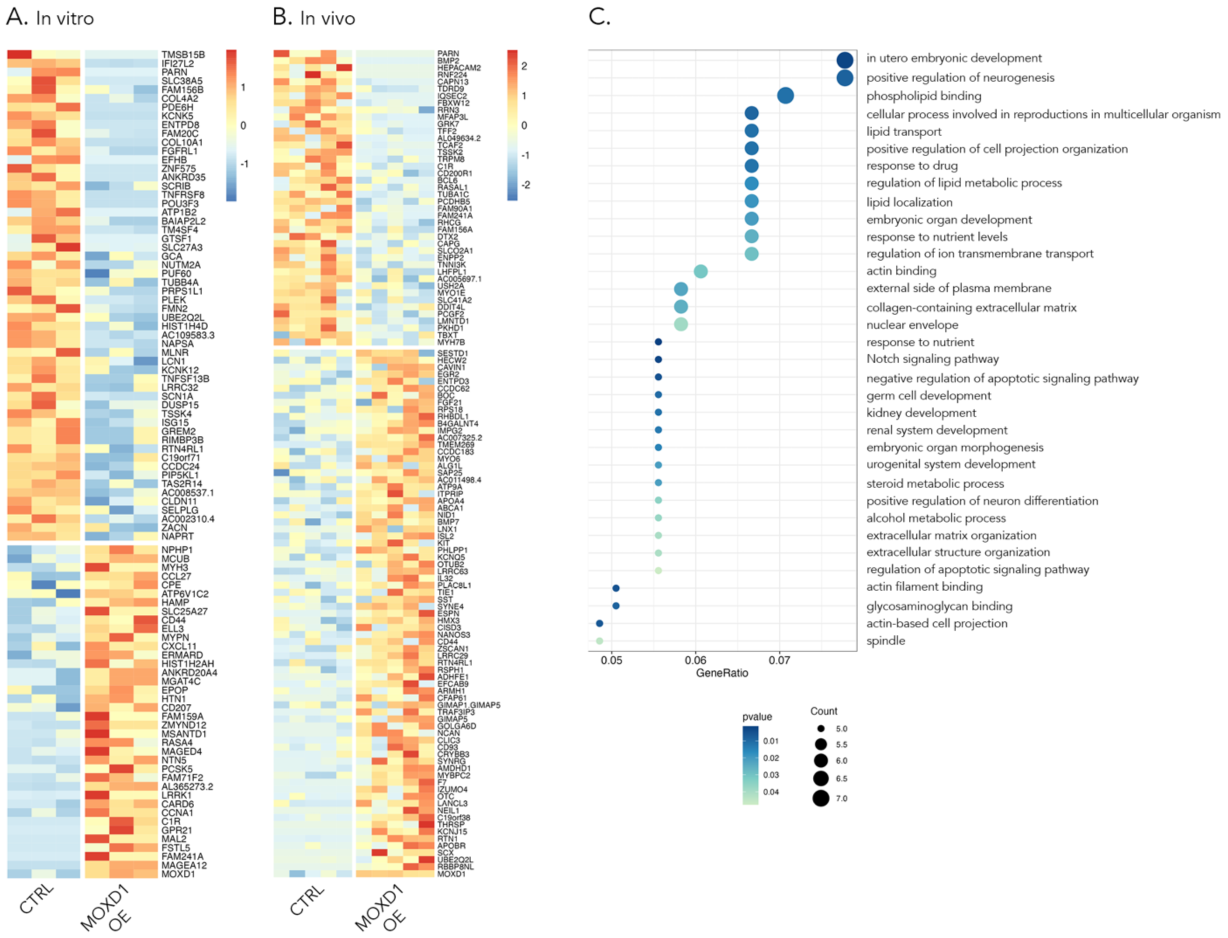

**Fig. S5. RNAseq of MOXD1 overexpressing NB cells *in vitro* and *in vivo* and Gene ontology enrichment analysis. (A-B)** Heatmaps of the identified DEGs between wild-type (CTRL) and *MOXD1* overexpressing (MOXD1 OE) SK-N-BE(2)c cells *in vitro* (A) and *in vivo* (B). **(C)** Gene ontology enrichment analysis of the DEGs in (B).

| IF antibodies | | | | |
|---|---|---|---|---|
| **Primary Antibody** | **Species** | **Dilution** | **Source** | **Product #** |
| HNK1 | Mouse | 1:5 | Hybridoma bank | 3H5 |
| MOXD1 (chick and human) | Rabbit | 1:1000 | Thermo Fisher | PA5-31526 |
| MOXD1 (zebrafish) | Rabbit | 1:200 | Aviva | ARP62346_P050 |
| **Secondary Antibody** | **Species** | **Dilution** | **Source** | |
| Anti-Mouse Alexa Fluor-594 | Goat | 1:1000 | Thermo Fisher | A-11032 |
| Anti-Rabbit Alexa Fluor-546 | Donkey | 1:1000 / 1:500 | Thermo Fisher | A-10040 |
| Anti-Rabbit Alexa Fluor-647 | Donkey | 1:1000 | Thermo Fisher | A31573 |
| Anti-Mouse Alexa Fluor-488 | Goat | 1:1000 | Thermo Fisher | A-11008 |

| IHC antibodies | | | | |
|---|---|---|---|---|
| **Primary Antibody** | **Species** | **Dilution** | **Source** | **Product #** |
| CD56/NCAM | Rabbit | 1:200 | Sigma-Aldrich | ab5032 |
| HNK1 | Mouse | 1:10 | DSHB | 3H5 |
| MOXD1 (chick) | Rabbit | 1:1000 | Thermo Fisher | PA5-31526 |
| MOXD1 (human embryos) | Rabbit | 1:100 | Thermo Fisher | PA5-31526 |
| MOXD1 (TMA) | Rabbit | 1:1000 | Thermo Fisher | PA5-31526 |
| phospho-Histone H3 (Ser10) | Rabbit | 1:200 | Sigma-Aldrich | 06-570 |
| RSPO3 | Rabbit | 1:200 | Abcam | 233113 |
| TH (chick) | Mouse | 1:10 | DSHB | ATH |
| TH (human embryos) | Rabbit | 1:100 | Abcam | ab112 |
| **Secondary Antibody** | **Species** | **Dilution** | **Source** | |
| anti-rabbit IgG (whole molecule)-Biotin antibody | Goat | 1:200 | Sigma-Aldrich | B-7389 |
| anti-mouse IgM (μ-chain specific)-Biotin antibody | Goat | 1:200 | Sigma-Aldrich | B-9265 |

| Nuclear staining | | | | |
|---|---|---|---|---|
| | **Species** | **Dilution** | **Source** | **Product #** |
| DAPI | | 1:3000 | Dako | D3571 |

**Table S1.** List of antibodies.

| **Target gene** | **5' - 3'** | |
|---|---|---|
| *UBC (Reference gene)* | Fwd | ATTTGGGTCGCGGTTCTTG |
| | Rev | TGCCTTGACATTCTCGATGGT |
| *SDHA (Reference gene)* | Fwd | TGGGAACAAGAGGGCATCTG |
| | Rev | CCACCACTGCATCAAATTCATG |
| *YWHAZ (Reference gene)* | Fwd | ACTTTTGGTACATTGTGGCTTCAA |
| | Rev | CCGCCAGGACAAACCAGTAT |
| DBH | Fwd | GCTCTCATGGAATGTCAGCTACA |
| | Rev | ACAGGACGCCAGCCTTGA |
| MOXD1 | Fwd | TGAGATGTTCCAAGACAGACAA |
| | Rev | TGCCACACACCAAAGGTTC |
| NOTCH1 | Fwd | CCGCAGTTGTGCTCCTGAA |
| | Rev | ACCTTGGCGGTCTCGTAGCT |
| PHOX2A | Fwd | GATGGACTACTCCTACCTCAATTCG |
| | Rev | GCTGCAGGCGCCAAAGT |
| PHOX2B | Fwd | CAGGGACCACCAGAGCAGT |
| | Rev | CTGCTTGCGCTTCTCGTTGA |
| PRRX1 | Fwd | CAGAACCGAAGAGCCAAGT |
| | Rev | GAGTAGGATTTGAGGAGGGAAG |
| SNAI2 | Fwd | ATGTCGGTTGTCTGGTTG |
| | Rev | TCTCCTGTGTTTTGTTCTTG |

**Table S2.** List of primers for qPCR.
